# Activation mechanism of the human Smoothened receptor

**DOI:** 10.1101/2022.06.07.493647

**Authors:** Prateek D. Bansal, Soumajit Dutta, Diwakar Shukla

## Abstract

Smoothened (SMO) is a membrane protein of the Class F subfamily of G-Protein Coupled Receptors (GPCRs) and maintains homeostasis of cellular differentiation. SMO undergoes conformational change during activation, transmitting the signal across the membrane, making it amenable to bind to its intracellular signaling partner. Receptor activation has been studied at length for Class A receptors, but the mechanism of Class F receptor activation remain unknown. Agonists and antagonists bound to SMO at sites in the Transmembrane Domain (TMD) and the Cysteine Rich Domain have been characterized, giving a static view of the various conformations SMO adopts. While the structures of the inactive and active SMO outline the residue-level transitions, a kinetic view of the overall activation process remains unexplored for Class F receptors. We describe SMO’s activation process in atomistic detail by performing 300 *μ*s of molecular dynamics simulations and combining it with Markov state model theory. A molecular switch, conserved across Class F and analogous to the activation-mediating D-R-Y motif in Class A receptors, is observed to break during activation. We also show that this transition occurs in a stage-wise movement of the transmembrane helices - TM6 first, followed by TM5. To see how modulators affect SMO activity, we simulated agonist and antagonist-bound SMO. We observed that agonist-bound SMO has an expanded hydrophobic tunnel in SMO’s core TMD, while antagonist-bound SMO shrinks this tunnel, further supporting the hypothesis that cholesterol travels through a tunnel inside Smoothened to activate it. In summary, this study elucidates the distinct activation mechanism of Class F GPCRs and shows that SMO’s activation process rearranges the core transmembrane domain to open a hydrophobic conduit for cholesterol transport.

## Introduction

G protein-coupled receptors (GPCRs) act as molecular telephones and transmit signals across the cellular membrane by associating with G proteins^1,2^ or arrestins.^3^ The process of signal transduction generally involves GPCRs binding to agonists which aid the shift in conformational equilibrium, facilitating the receptors to transition to an *active* state. Activation allows the receptor to associate with intracellular binding partners, allowing the process of signal transduction.^4^ GPCR activation is an area of active research - with studies establishing conserved structural motifs like the E/DRY, NPxxY^4–8^ in Class A and PxxG, HETx^9^ in Class B receptors acting as molecular switches that stabilize the inactive state. Unlike Class A and B GPCRs, activation of Class F receptors : Smoothened (SMO), Frizzleds1-10 (FZD_1–10_) is still poorly understood. A primary reason for this elusiveness is that these receptors share none of the structural motifs seen in Class A/B, and have less than 10% sequence similarity to Class A receptors^10^ as well as Class B receptors. Since Class A and B GPCRs are involved in mediating virtually every physiological response : they are crucial drug targets, as 34% of all FDA approved drugs target one of these proteins.^11^

Smoothened (SMO) is a transmembrane protein from the Class F of GPCRs. Class F consists of proteins that are involved in maintaining tissue homeostasis and regenerative responses in adults, and are crucial in embryonic development, as they regulate cellular differentiation by binding to sterol and Wnt ligands.^12–15^ SMO is expressed in tissues throughout the body, particularly in cerebellar and pituitary tissue,^16^ and is a member of the Hedgehog (HH) signaling pathway. When the endogenous inhibitor of SMO, a membrane protein Patched (PTCH), is inhibited by Sonic Hedgehog (Shh) binding, SMO translocates to the ciliary membrane, and undergoes conformational transitions (activation) to bind to its intracellular signaling partner G_*i*_.^17,18^ How PTCH inhibits SMO is still unclear. However, multiple studies have described PTCH’s inhibition on SMO as acting through reducing SMO’s accessibility to membrane cholesterol.^19^ A recent study described the effect of PTCH on the cholesterol accessibility of the upper leaflet, suggesting that PTCH inhibits SMO by either transporting cholesterol to the inner leaflet, or to an extracellular acceptor.^20^ HH signaling is critical to embryonic development, and any changes in signaling can lead to severe birth defects.^21^ Cyclopamine, a naturally occuring alkaloid in corn lily, has been identified as a teratogen (agents responsible for birth defects in infants),^22^ and was responsible for birth defects in lambs in Idaho in the 1950s.^23^ It was identified later that cyclopamine’s mechanism of action involved inhibiting HH signaling by binding to SMO.^24–26^ On the other hand, overstimulation of HH signaling via SMO has been linked to the pathogenesis of pediatric medulloblastoma and basal cell carcinoma.^27,28^ Vismodegib^29^ and Sonidegib^30^ are two FDA approved drugs that target SMO, but are prone to chemoresistance. ^31^ Therefore, understanding activation mechanisms of Class F GPCRs is hence critical to design novel therapeutics.

Structures of SMO bound to agonists and antagonists outline the effects of allosteric and orthosteric modulators binding on SMO activity. These structures show the existence of two primary binding sites in SMO - the first in the Cysteine Rich Domain (CRD), which binds agonists cholesterol^32^ and cyclopamine. ^34^ The second site is present in the TMD, which binds both antagonists LY2940680,^10^ SANT1 and AntaXV,^35^ cyclopamine,^36^ TC114,^37^ Vismodegib^32^ and agonists SAG1.5,^35^ SAG,^38^ SAG21k,^39^ 24, 25-epoxycholesterol,^33^ and cholesterol. ^38^ Mutagenesis studies have outlined the presence of an intracellular W^7.55*f*^-R^6.32*f*^ *π*-cation lock^40^ in Class F that is broken on activation (Fig. 1A), with mutations that disrupt this lock lead to increased agonist potency and pathway selection (superscripts refer to the modified Ballesteros-Weinstein numbering system used to denote Class F GPCR TM residues^41^ introduced by Wang et al.^35^). On the extracellular end, for SAG-bound SMO, the D-R-E network is broken in active SMO^35^ (Fig. 1A). The intracellular end of active SMO shows rearrangements in TM6 (outward), TM3 (outward) and TM5 (inward) (Fig. 1B). These studies paint a static picture of how SMO activity can be attributed to structural rearrangements, however, a dynamic understanding of the process of SMO activation still remains. Hence to provide a dynamic overview of activation, we simulated ~ 250 *μ*s Apo-SMO (no ligand bound) to understand SMO’s activation process in atomistic detail. Moreover, it has been shown that PTCH modulates SMO activity by controlling its access to membrane cholesterol^19,42^ which then travels through a hydrophobic tunnel inside SMO to access the primary ligand binding site in CRD, showing an expanded tunnel in active SMO (Fig. 1A). Hence, we simulated agonist bound (SAG-SMO) (~ 36 *μ*s) and antagonist bound (SANT1-SMO) (~ 42 *μ*s to explore the effects of bound modulators on SMO activity, and the mechanisms of action for these molecules. Using a highly parallel Adaptive sampling based approach and constructing a Markov state model (MSM),^43,44^ we probe submillisecond dynamics of SMO, and show that SMO activation involves a intracellular structural motif that is conserved across Class F receptors. MSMs have been used to model membrane protein behavior at varied timescales, to probe activity of membrane transporters,^45–47^ as well as to study conformational dynamics of signaling proteins.^8,48–55^ In particular, Markov state models have been employed to investigate conformational dynamics of GPCRs, such as beta2-adrenergic receptor,^8,52,56^ Dopamine D_3_ receptor,^48^ *μ*-opioid receptor, ^54,55^ Chemokine receptor CCR2,^49^ and Cannabinoid Receptors 1, 2.^50,51^ Using MSMs, we outline the involvement of multiple CRD-TMD salt-bridges that are rearranged during SMO activation, establishing a role for the CRD in SMO activation. We show that the hydrophobic tunnel inside SMO expands in the presence of an agonist, and is occluded by the antagonist. These observations are amenable to experimental observations that bolster the cholesterol transport-like activity of SMO. We then use a mutual-information based approach to outline the allosteric mechanisms through which the agonist SAG operates, i.e. by changing the allosteric pathways in SMO to more active-like SMO. These observations provide a detailed and atomistic in-depth view of SMO activation, and may aid in design of antagonists for cancer therapy.

**Figure 1:**
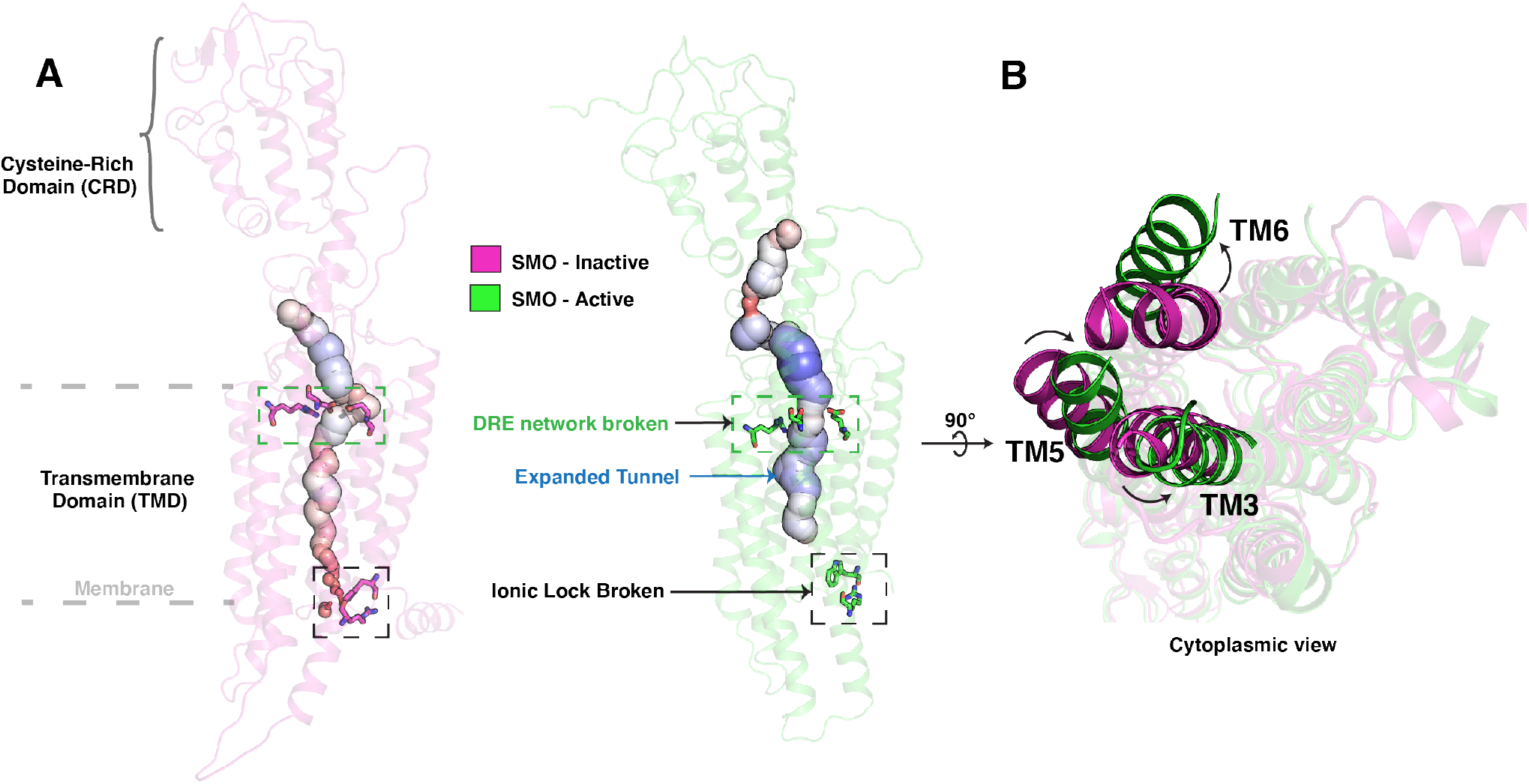
Major structural changes during SMO Activation. (A) Comparison of the broken D-R-E network and the W-R *π*-cation lock, and the expanded tunnel, in inactive (magenta, 5L7D^32^) vs active (green, 6XBL^33^) SMO (B) Comparison of inactive and active SMO, indicating the outward movement of the TM6 and TM3 and inward movement of TM5 in active SMO.

## Results and Discussion

### SMO activation involves a conserved molecular switch

To probe the transitions SMO undergoes during activation, SMO was simulated in a ligand-free form (Apo-SMO) from two starting points - inactive and active structures. Simulations were performed using a parallel approach - by clustering the existing data based on selected features (feature selection explained in Methods) and seeding the next round of simulations by randomly selecting starting points from the least populated clusters - a technique known as Adaptive sampling^57^ (Fig. S8, Table S1, S2). The high dimensionality of the data was reduced by transforming it using time-Independent Component Analysis (tICA). ^58,59^ tICA uses a linear combination of the supplied features to identify the slowest collective degress of freedom in the data by computing the time-lagged autocorrelation. The first two tICA components account for the two slowest processes associated with activation (Fig. S9, S10). The active and inactive structures were separated majorly in the first tICA component (tIC 1), indicating that activation was the slowest process observed in simulations. Hence, features that were highly correlated with tIC 1 (Fig. S11) were considered pivotal to activation. The convergence of the data, clusters and hence the free energies derived from it, were confirmed by the presence of a continuous density of data along tIC 1 (Fig. S9A). This shows that the simulations have indeed sampled the conformational landscape necessary to probe the activation pathway of SMO. The tICA transformed data was clustered - dividing the data into kinetically distinct microstates. A MSM was constructed on the clustered data to compute the transition rates between microstates, and to reweigh the data, eliminating the bias introduced by Adaptive sampling.

At the intracellular end, we observe that W339^3.50*f*^ shows a very dramatic reorientation on receptor activation, moving outwards from the center of the TM bundle, to accommodate the bound G_*i*_. W339^3.50*f*^ is conserved across all Class F receptors (Fig. S12). Upon further analysis, we ob that this rearrangement extends to include M449^6.30*f*^ and G453^6.34*f*^ (outward movement), G422^5.65*f*^ (translation) (Fig. 2A), as well as W535^7.55*f*^ (inward rotation) - residues that are all conserved across the entire Class F family (Fig. S12).

**Figure 2:**
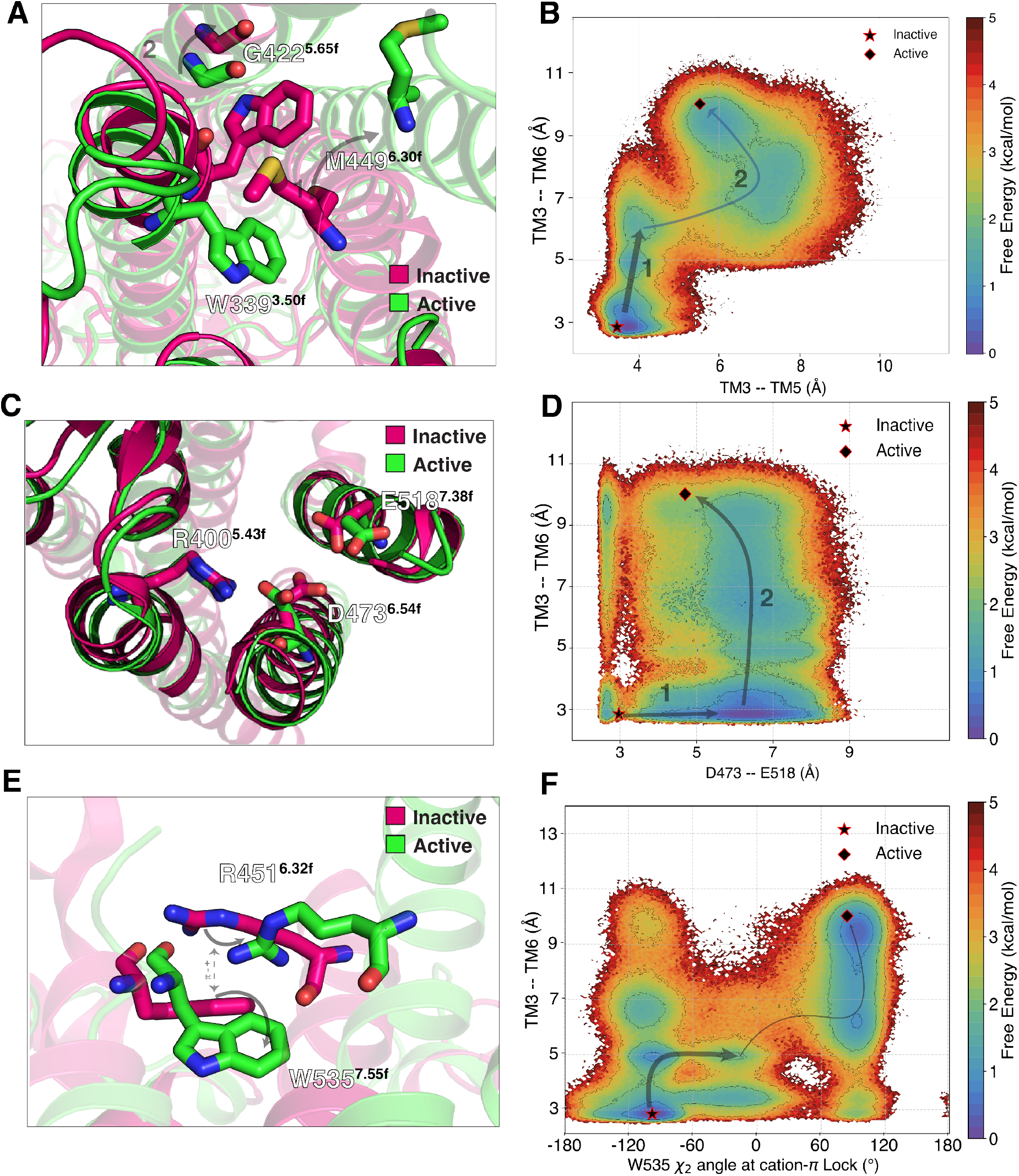
Molecular metrics integral to SMO Activation. (A) Rearrangement of the WGM motif, a conserved molecular switch across class F GPCRs, undergoes rearrangement on SMO activation. (B) Relative free energies from MSM-weighted simulation data plotted on the TM3-TM6 distance vs TM3-TM5 distance measured at residues W339^3.50*f*^, M449^6.30*f*^ and G422^5.65*f*^. (C) Breaking of the D-R-E network on the extracellular end of the TMD. (D) Similar to (B), but for TM3-TM6 distance vs the D-E distance. (E) The *π*-cation lock breaks by the sidechain rotation of W535^7.55*f*^. (F) Same as (B) but for TM3-TM6 distance vs χ_2_ dihedral measured at W535^7.55*f*^.

M449^6.30*f*^’s outward movement is a proxy for the outward movement of TM6 a process associated with canonical GPCR activation.^33,38^ However, instead of kinking outwards as observed in Class B receptors, TM6 in SMO undergoes translation, to accommodate G_*i*_. This can be attributed to the absence of P^6.43*f*^, a residue conserved across FZDs (Fig. S12). P^6.43*f*^ is replaced by F462^6.43*f*^ in SMO - thereby increasing its rigidity and resistance to developing kinks.^60^ Recently published structure of active FZD7^61^ shows this kink at P^6.43*f*^. Similar translation in TM6 is observed in Rhodopsin, a Class A receptor. ^62^ This particular feature is hence unique to the activation mechanism of SMO. TM5 on the other hand, shows slight inward translation. To capture these outlined movements, we projected the entire Apo-SMO data onto W339^3.50*f*^ – M449^6.30*f*^ (TM3-TM6 distance) v/s W339^3.50*f*^ – G422^5.65*f*^ (TM3-TM5 distance) and computed the free energy associated with each state(Fig. 2B, Fig. S13). The free energy plot shows that this TM3-5-6 rearrangement follows a stage-wise process - with TM6 moving outwards first by ~ 4 Å, (State **1** in Fig. 2B) followed by the rest of the TM3 outward movement after a slight outward rearrangement in TM5 (State **2** in Fig. 2B). The overall free energy barrier for this rearrangement is ~ 2.5 kcal/mol. The outward movement of TM6 is analogous to class A receptor activation (Fig. S14 A,B). A conserved molecular switch mediating SMO’s activation on the intracellular end is similar to the breakage of molecular switch E/DRY in Class A GPCRs-with W339^3.50*f*^ being the residue analogous to R^3.50^ (Fig. S14 C,D). Hence, we posit that this conserved molecular motif (W-G-M) is integral to Class F receptor activation, and provides a basis for activation across the entire Class F receptors, while also showing the uniqueness of activation of Class F receptors.

The crystal structure of SMO bound to the synthetic agonist SAG1.5 gives clues about the activation-specific residue-level rearrangements that occur on the extracellular end of SMO. D473^6.54*f*^ has been established as a residue critical to SMO activity, as it forms a part of SMO’s core TMD ligand binding cavity, and is shown to interact with agonists SAG1.5, SAG, oxysterols and antagonists GDC-0449, AntaXV.^33,35,38,63,64^ Specifically, a network of salt bridges formed by the residues D473^6.54*f*^, E518^7.38*f*^ and R400^5.43*f*^ is broken in SAG1.5-bound SMO (Fig. 2C).^35^ Hence, we also projected the Apo-SMO data on the D473^6.54*f*^ – E518^7.38*f*^ distance v/s intracellular TM3-6 movement (Fig. 2D, Fig. S13). We observe that the TM6-TM3 outward movement (**2** in Fig. 2D) is preceded by the breakage of the hydrogen bond between D473^6.54*f*^-E518^7.38*f*^ (**1** in Fig. 2D).

To outline the role of the *π*-cation lock W535^7.55*f*^-R451^6.32*f*^ in activation, we projected this *π*-cation lock contact v/s the TM3-6 outward movement (Fig. 2F, Fig. S13) for Apo-SMO. Projecting the Apo-SMO data along the sidechain dihedral angle χ_2_ of W535^7.55*f*^, clearly showed the distinct inactive and active states. This shows that the mechanism of *π*-cation lock breaking involves the sidechain rotation of W535^7.55*f*^. Additionally, we observe that the *π*-cation lock breaks around the same TM3-TM6 distance as the outward movement of TM3. Thus, the WGM motif and the *π*-cation lock at the intracellular end, and the D-R-E network at extracellular end are critical residue networks involved in SMO activation. These residues form a network of allosterically coupled residues, proving crucial for signal transduction across the membrane.

### Residues at the CRD-TMD interface involve salt-bridge rearrangements in SMO activation

SMO, in addition to a heptahelical TM domain, possesses an extracellular domain called the Cysteine Rich Domain (CRD). The CRD consists of residues that are highly polar in comparison to the TMD, which is mostly hydrophobic (Fig. S15). This domain is critical for SMO activation, as SMOΔCRD mutants show a higher constitutive activity - suggesting that the CRD represses SMO’s basal activity.^65^ The CRD also includes the primary sterol binding site in SMO^32^ - and it has been posited that PTCH inhibits SMO by reducing cholesterol access to this site. ^18^ Structures of active xenopus laevis SMO (xSMO) show a dramatic reorientation of the CRD on xSMO activation-suggesting that the CRD has a very dynamic range of motion. ^64^ However, this reorientation is not observed in human SMO (hSMO). ^32,37^ Thus to establish a role of the CRD in activation of hSMO, we sought residue pairs in Apo-SMO CRD-TMD interface that showed the highest variance along tIC1, the slowest process that captured Apo-SMO activation.

Fig. 3(A-F) show the residue pairs that have the highest change in contact frequency during activation - starting with the R485^6.66*f*^ – D209^*CRD*^, salt-bridge, which breaks during activation (Fig. 3A) due to the outward movement of TM6. This indicates that the R485^6.66*f*^–D209^*CRD*^ salt-bridge is involved in stabilizing the inward conformation of TM6 in the inactive state. This loss of the R485^6.66*f*^–D209^*CRD*^ salt-bridge is however compensated by the formation of the nearby R161^*CRD*^–D486^6.67*f*^ salt bridge, which is predominantly seen in the active conformation (Fig. 3E). Furthermore, the inactive state shows a salt-bridge E208^*CRD*^–K395^*ECL*2^ which breaks on activation, compensated by the formation of the nearby D201^*CRD*^–R296^*ECL*1^ (Fig. 3B, E). Additionally, activation strongly favors the formation of R159^*CRD*^–D209^*CRD*^ (Fig. 3C) and D382^*ECL*2^-K204^*CRD*^ (Fig. 3G) salt bridges. The inactive (green) and active (magenta) structures depicted in the figure were taken as representative structures from the inactive-like and active-like free energy wells in the tIC landscape.

**Figure 3:**
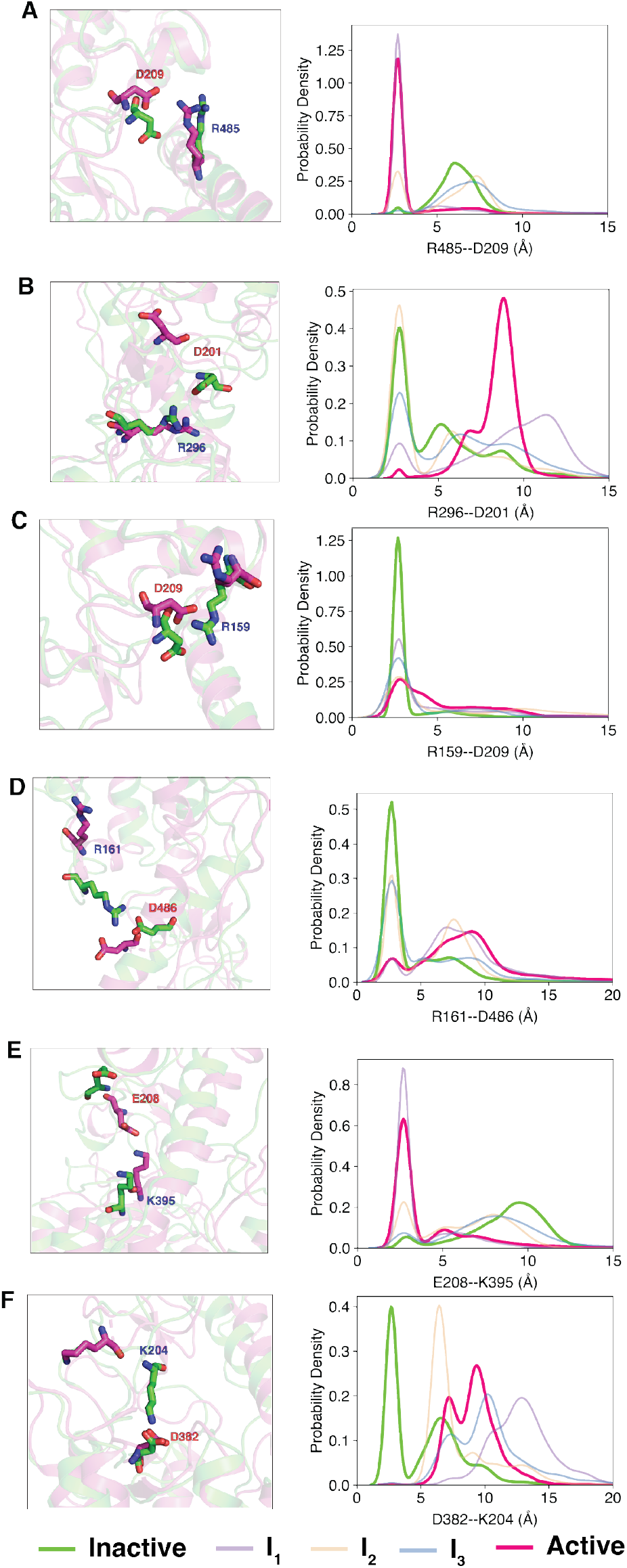
Overall activation of SMO involves residues at CRD-TMD junction. (A)-(F) Snapshots and probability density plots outlining the salt-bridge rearrangements at the CRD-TMD interface during SMO activation.

The path along tIC1 from the inactive state to active state involves 3 intermediate states(I_1–3_) (Fig. 4A), characterized by free energy barriers of atleast 1 kcal mol^−1^ among them. Using Transition Path Theory on the constructed MSM, we calculated the fluxes of transitions between these states, to establish timescales for activation of SMO (Fig. 4B). The simulations show that the entire process of activation from inactive to active has a MFPT (mean first passage time) of ~ 72*μ*s (Fig. 4B), while the reverse process is ~ 3X faster, with MFPT ~24 *μ*s.

**Figure 4:**
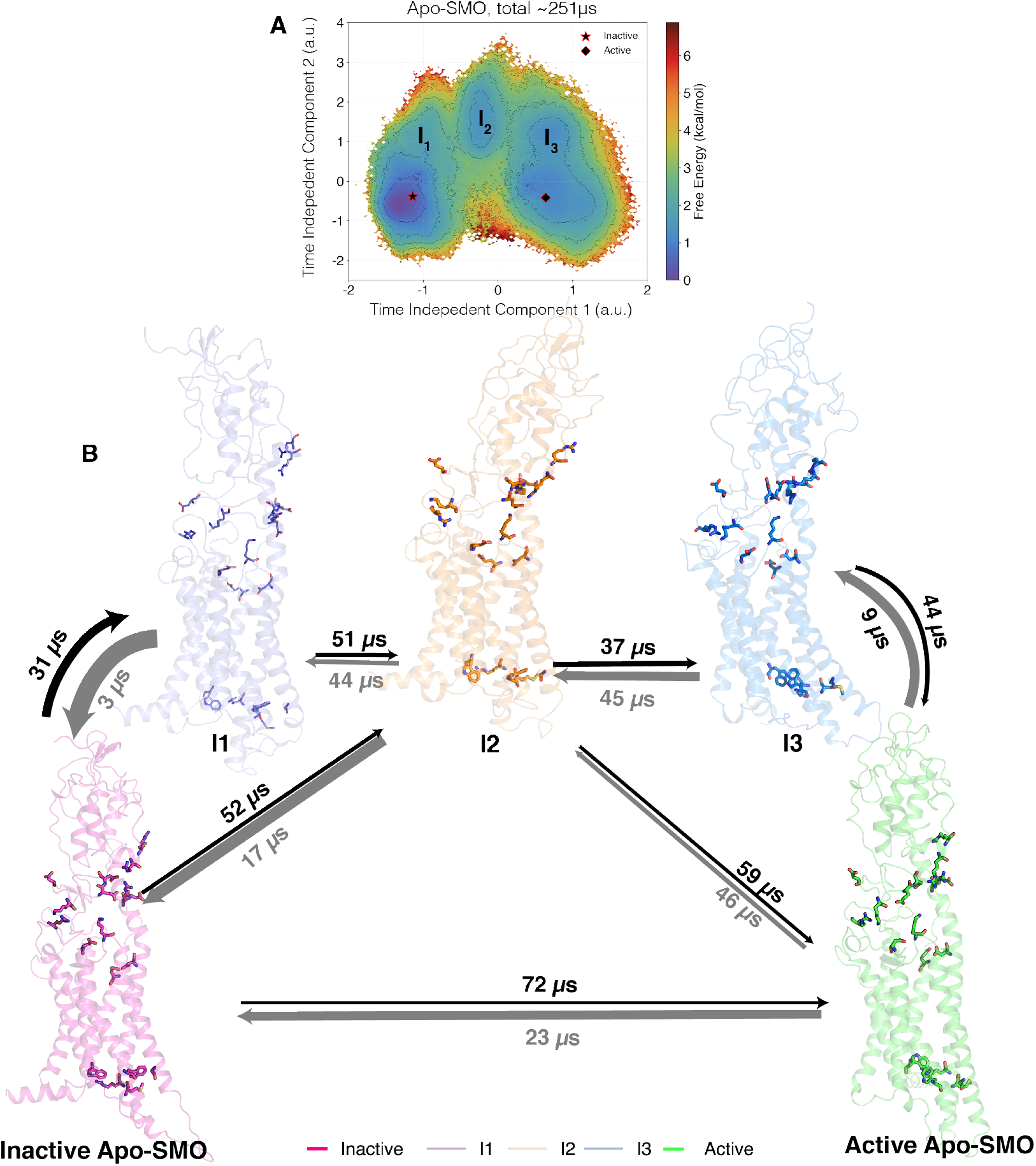
(A)Relative free energies from MSM-weighted simulation data of Apo-SMO plotted along tIC1 and tIC2, the 2 slowest components, with the intermediate states I_1–3_ as shown. The intermediate states I_1–3_ were defined based on metastable basins and free energy barriers associated with transitioning from an inactive to an active state. A cutoff of 1.8 kcal/mol was used to separate one basin from another. Residues shows as sticks include the *π*-cation lock, the WGM motif and the salt bridges involved in activation. (B) Overall transition pathway of SMO activation process. The inactive (PDB ID: 5L7D)^32^ and active (PDB ID: 6XBL)^38^ structures are separated by the presence of 3 metastable conformations in between, I_1–3_. Residues shown by sticks correspond to the salt bridges, the WGM motif, the DRE network and the *π*-cation lock, all residues critical for mediating SMO activation.

We observe that residue pair rearrangements that are associated with activation at the CRD-TMD junctions are salt-bridges, mostly between residues with one residue in CRD and the other one in TMD (Fig. S16). Almost none of these polar residues are conserved (Fig. S12, S17), indicating that these residues contribute to a unique activation process for SMO at the CRD-TMD interface. Additionally, we observe that the entire CRD motion can be accounted for by a slight outward rotational motion of the CRD (Fig. S18), thereby causing TM6 to move outwards and triggering activation on the intracellular end. Since the CRD has a cholesterol binding site, it is possible that cholesterol binding to the CRD triggers this outward rotation, inducing the signal that causes TM6 to move out. This potentially outlines a mechanism for the activation of SMO by cholesterol, its endogenous agonist.

### SMO’s Activation is linked to opening of a hydrophobic tunnel

Endogenously, on PTCH’s inhibition by Shh, SMO is activated. SMO’s activation is mediated endogenously by cholesterol, suggesting that PTCH’s inhibition facilitates SMO’s activation by cholesterol. This suggests that cholesterol from the membrane travels to the extracellular sterol binding site. How this transfer of cholesterol occurs to the SMO CRD is still unknown. However, SMO does indeed present itself with a unique topology - the presence of a tunnel inside the protein. This tunnel has been hypothesized^33,38,39,64^ to facilitate the transport of cholesterol from the membrane to the binding site in the CRD,^32^ making this tunnel a prime target for inhibitors. As noted by Qi et al., SMO antagonists (SANT1, AntaXV, LY2940680) bind deeper into a tunnel inside SMO, whereas SMO agonists (SAG) bind outside this tunnel. Adding a 4-aminomethyl moiety to the tail-end of SAG converts it to an antagonist, suggesting that this added moiety can hinder the tunnel. ^66^ Mutations that introduced a bulky residue into the tunnel (V329F, V333F, V408F,I412F,T470Q), blocked SMO activity, ^32,39^ suggesting that the tunnel conformation was linked to how small molecule and mutations modulated SMO activity.^38^ This suggests that SMO antagonists like SANT1 act as steric antagonists by blocking the sterol tunnel inside SMO, while agonists like SAG allosterically activate SMO by breaking the D-R-E network, setting off receptor activation on the intracellular end. The mechanism and dynamics of the modulators acting on SMO’s activation is still unclear. Hence we simulated SMO bound to antagonist SANT1 (SANT1-SMO) and agonist SAG (SAG-SMO) to probe the effect of bound agonist and antagonist on SMO’s activation.

SMO’s tunnel is characterized by markedly hydrophobic residues (Fig. S19), pointing further towards the idea that a hydrophobic molecule may be transported through it. This tunnel runs through the core of the receptor, spans the entire TM domain, starting at the conserved residues W339^3.50*f*^, spans ~ seven helical turns, and ends at the extracellular network of residues E518^7.38*f*^, D473^6.54*f*^ and R400^5.43*f*^. These three residues form the base of the space between the CRD and TMD. Moving outwards along the path defined along the tunnel directly leads to the binding site, with TM6, ECL2 and ECL1 forming the bridge between these sites (Fig. S20).

In SANT1-SMO simulations, the tunnel remains almost completely blocked (Fig. 5A, B), indicating that the mechanism by which SANT1 modulates SMO activity is by binding deeply into the SMO tunnel core, precluding the potential transport of cholesterol. SANT1’s piperazine moiety directly interacts with H470^6.51*f*^ and sidechain of M525^7.45*f*^ - forming hydrogen bond interactions. The pyrrolic head of the ligand remains buried deep inside, with minimal movement normal to the plane of the membrane, along the tunnel (Fig. S21). However, in Apo-SMO simulations, the tunnel remains relatively open(Fig. 5C, D). Interestingly, we observe a conformational dependence of the lipid organization in the membrane - Inactive SMO surrounds itself with a cholesterol in the upper leaflet, as opposed to other cases (Fig. S22). This suggests that cholesterol shows a propensity to accumulate outside inactive SMO to possibly transport itself in the hydrophobic tunnel, leading to SMO activation. Additionally, In SAG-SMO simulations, we observe that the tunnel radius has a sudden kink outward (z ~ −20 Å), suggesting that there is a relative expansion of the tunnel induced by SAG (Fig. 5 E, F). In the simulations, the membrane extends from z = 0 to z = −40. Since this expansion occurs between z = 0 and z ~ −20 Å, it suggests the opening is in the upper leaflet (Fig. S24A). On plotting the free energy difference between Apo-SMO and SAG-SMO, a marked difference in the free energy associated with the opening of the tunnel is observed (Fig. S23A-C). Recent studies suggest that active PTCH precludes SMO’s accessibility to cholesterol in the upper leaflet. ^20^ To further probe into the exact position of this tunnel opening, we observed that a cluster of openings occured at x 16 Å and y ~ 22 Å - corresponding to the space between TM2 and TM3 (Fig. S24B). This is in agreement with a recent study that used coarse-grained simulations to observe a cholesterol binding site at the TM2-TM3 interface in the upper leaflet. ^68^ Thus, SAG acts as an agonist by allosterically expanding the tunnel at the cholesterol interaction site - giving further evidence for the cholesterol-transport like activity of SMO. Thus we conclude that SANT1 functions as a steric antagonist by blocking the tunnel, whereas SAG functions by allosterically expanding the tunnel, thereby establishing design rules for SMO agonists and antagonists.

**Figure 5:**
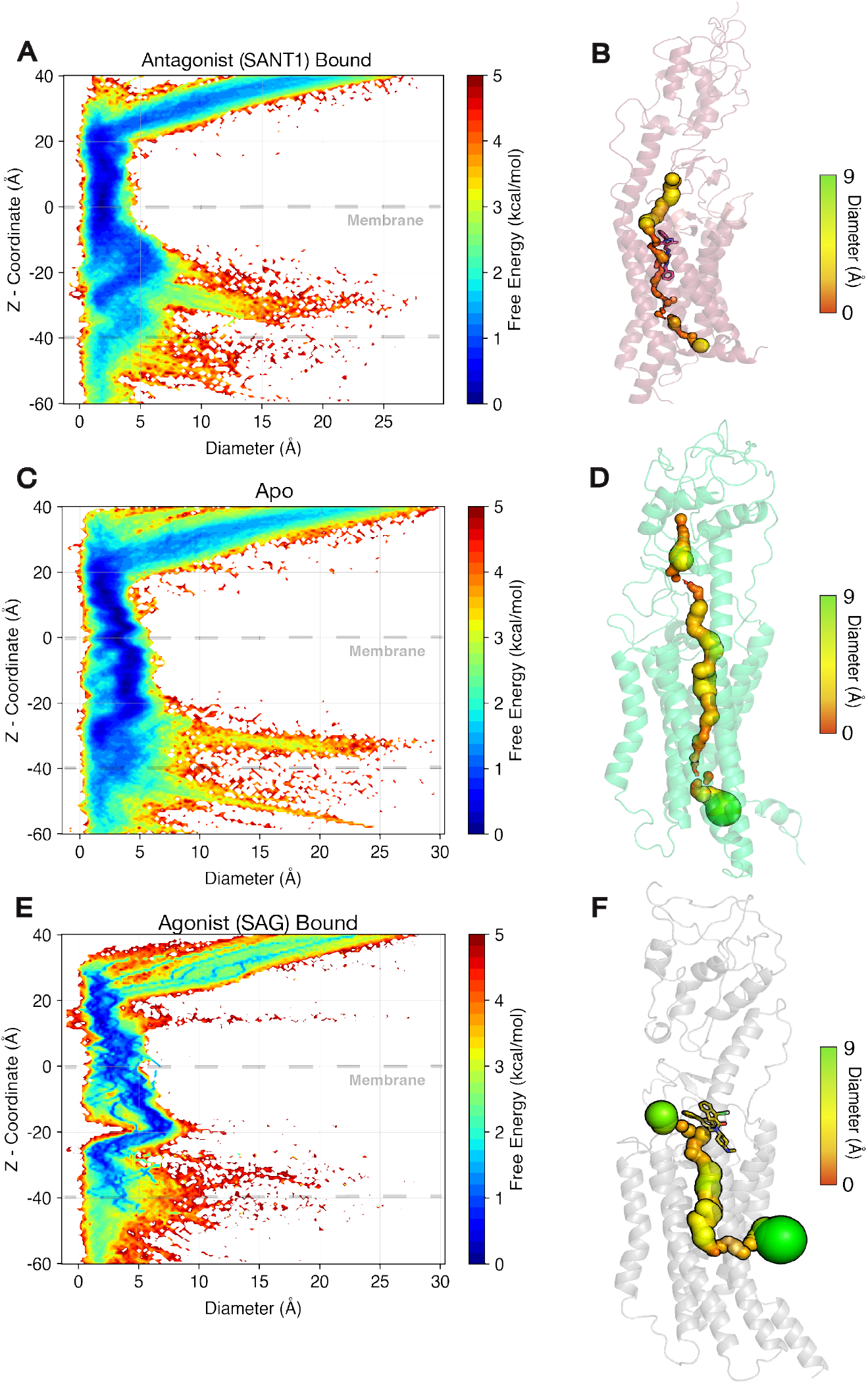
Tunnel radius plots for SMO. (A) Free energy plot of the tunnel diameter along the z-coordinate for SANT1-bound SMO. (C) same as (A), but for Apo-SMO. (E) same as (A), but for SAG-bound SMO. SAG-bound SMO clearly shows the expansion of the tunnel as compared to Apo-SMO and SANT1-SMO. (B), (D), (F) - representative figures for SANT-1 SMO, Apo-SMO and SAG-SMO. Tunnel radii were calculated using the HOLE program^67^

**Figure 6:**
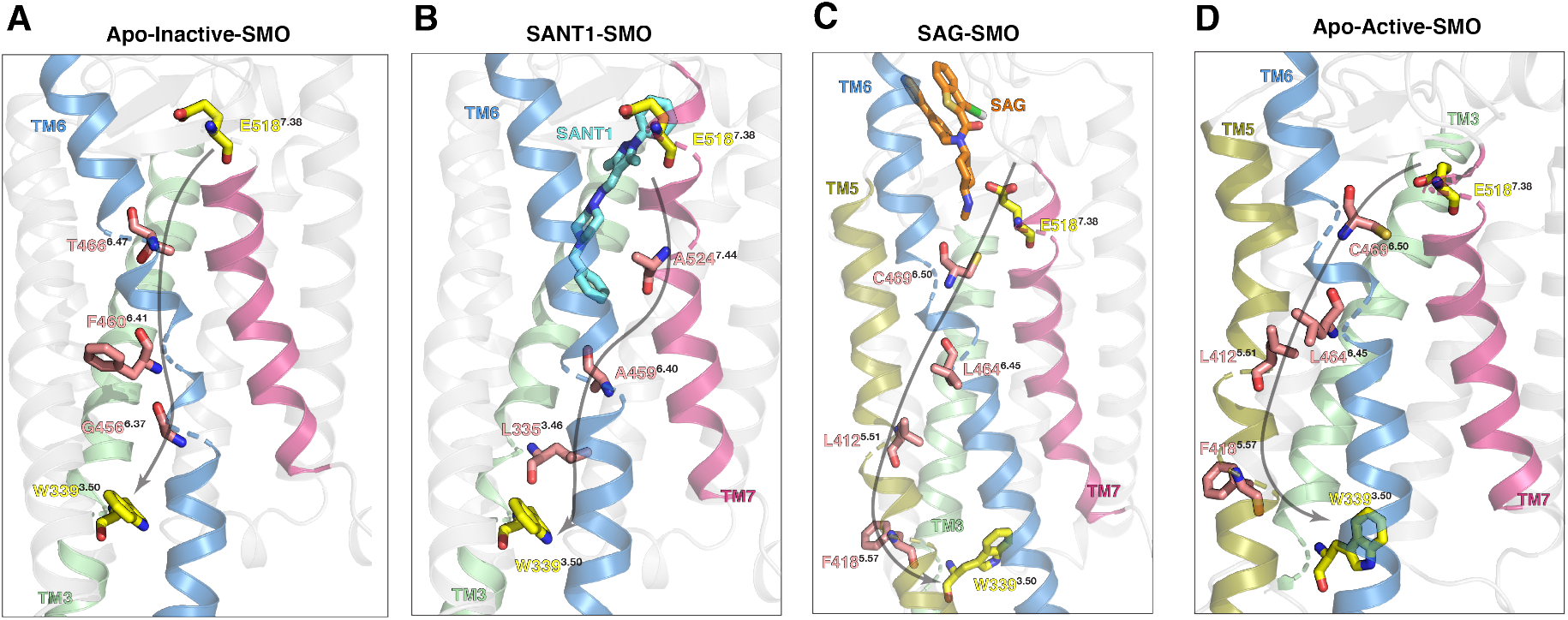
Allosteric pathways between E518^7.38*f*^ and W339^3.50*f*^. (A) Pathway in Apo-Inactive-SMO. Since the tunnel radius is decreased, TM6 outward movement is restricted, and therefore the entire allosteric communications occurs via TM6. (B) In SANT1-SMO, due to slight outward movement of TM6, the pathways switches from TM7 to TM6 to TM3. (C,D) SAG-SMO and Apo-Active SMO show the same allosteric pathway, which spans TM7-TM6-TM5-TM3.

### SAG alters the allosteric pathways in SMO during the process of SMO activation

To further investigate the mechanism by which SAG allosterically modulates SMO’s activity resulting in the expansion of the tunnel, we computed the allosteric pathways that connected the intra- and extracellular ends of SMO, responsible for transmembrane signal transduction. Allosteric pathways contain a series of conformationally-coupled residues that link dynamically active and spatially distant residues. In Class A GPCRs, allosteric pathways are responsible for communicating conformational changes from the extracellular end to the intracellular end, completing the process of signal transduction.^69–71^ Since SMO’s activation process involves allosteric communication between the extracellular ligand binding site (D-R-E network) and the G-protein coupling site (WGM motif), we sought to analyze the allosteric pathways that connect the two sites. We computed the dynamic pairwise mutual information of Inactive-Apo-SMO, Active-Apo-SMO, SANT1-SMO and SAG-SMO on a residue-level basis, and constructed a graphical network of residues that are allosterically linked. The dynamic mutual information takes into account the residue-level movements Based on this network, we present the allosteric pathway between the intra- and extracellular ends of TMD.

In our simulations, we observe that the allosteric pathway between the intra and extracellular ends in Apo-Inactive SMO completely passes through TM6, encompassing residues T466^6.47*f*^, F460^6.41*f*^ and G456^6.37*f*^ (Fig. 5A). This establishes an integral role for TM6 in mediating the signals across the transmembrane domain in inactive-SMO. SANT1-SMO on the other hand, unexpectedly shows a distinct pathway, first going down intra-helically to A524^7.24*f*^, crossing over to TM6 via A459^6.40*f*^ and finally to L335^3.46*f*^. This however can be explained by the observation that the SANT1 causes a slight outward movement of TM6, to accommodate itself in the deep core TMD ligand binding cavity (Fig. S25). This outward movement of TM6 moves T466^4.67*f*^ away from E518^7.55*f*^. This causes the network to rearrange itself, moving over to TM6 further downstream. On the other hand, SAG and Apo-Active SMO show the exact same networks, further indicating that SAG alters the allosteric networks in SMO to resemble Apo-SMO. These networks involve C469^6.50*f*^, the most conserved residue in TM6, down L464^6.45*f*^, and a flip over to TM5 as we move intracellularly, due to the outward intracellular movement of TM6, via L412^5.55*f*^ and F418^5.61*f*^. Thus, we can establish a basis for the mechanisms through which SAG and SANT1 effectively modulate SMO activity, and establish an integral role for TMs 7,6,5,3 in signal transduction.

## Conclusions

Our study reveals the activation mechanism for SMO, a Class F GPCR, in atomistic detail via molecular dynamics simulations. We characterized the residue level transitions that SMO undergoes during activation. We simulated SMO in Apo, SAG and SANT1 bound states to probe the activation mechanism of SMO, and computed the free energy landscape of the process. Our MSM weighted free energy landscapes show a barrier of max free energy barrier of ~ 3 kcal mol^−1^ while transitioning from an inactive to active state, involving three intermediate states.

Class A and Class B receptors have been the subject of major interest involving GPCR activation. ^4,9^ Receptor activation studies on Class F majorly focused on the start and end states of the receptor, without giving an overview of the dynamics of the process. Using computational methods, we show that SMO activation involves the rearrangement of a intracellular structural motif : the W-G-M motif, conserved across the entire Class F family. This lays the basis for a common activation mechanism for all Class F receptors on the intracellular end. Additionally, this motif involves W^3.50*f*^, which is the residue equivalent to R^3.50^ in class A receptors, establishing the integral role of TM3 in GPCR activation. On the extracellular end of TMD, we see that the D-R-E network of residues is pivotal to activation, as it engages the agonist and sets off the activation process at the intracellular end. We also show evidence of allosteric coupling between these two sites, showing that the rearrangement of the D-R-E network is necessary to ensure intracellular rearrangement of the WGM motif.

We also establish a role for the CRD in SMO activation, forming and breaking salt-bridges while transitioning to an active state, contacts that have not been discussed previously. This gives novelty to the methodology established, inferring that MD simulations can be used to discover contacts crucial to activation, previously unknown. We show that the agonist SAG expands an intra-TMD tunnel inside SMO, further supporting the hypothesis that SMO transports a cholesterol molecule through its hydrophobic tunnel to activate SMO. ^19,20,33,38,64^ We also show that SAG acts as an allosteric modulator, by modifying SMO’s allosteric pathways to be similar to Apo-SMO. On the other hand, SANT1 acts as a steric antagonist, by occluding the hydrophobic tunnel inside SMO, hence lowering the radius. Therefore, we establish the mechanisms of action of antagonists and agonists in modulating SMO activity. Additionally, experimental validation by mutagenesis of the role of various residues needs to be performed for further corroboration of this computational study. Mutation of residues of the WGM motif - (W339, G422, M449) the various salt bridges, the interface of upper leaflet and TM2-3, and the allosterically coupled residues, possibly through techniques like Alanine scans and Deep Mutagenesis can be performed as testable hypotheses, thereby delineating the role of these residues in modulating SMO activity. Additionally, how cholesterol, the endogenous agonist of SMO, modulates SMO activity in the presence of agonists, still needs to be explored. However, we propose that the overall mechanistic findings from this study can be used to design novel SMO antagonists, for chemotherapy.

## Methods

### Molecular Dynamics (MD) Simulations

#### Simulation setup

SMO structures in the bound inactive conformation (inactive-SMO) (PDB ID: 5L7D^32^) and active conformation (active-SMO) (PDB ID: 6XBL^38^) were used as starting points for the SMO-Apo simulations. For apo systems, the bound ligand and the stabilizing antibodies were removed. The missing residues in the proteins were modeled using MODELLER^72^ (Table S3). The inactivating mutation V329F in the inac-SMO was corrected back to wild-type. For SAG-SMO, the bound SAG was retained in the SMO-SAG complex. ^38^ To check for the protonations in acidic residues under physiological conditions, the pKa was calculated using the H++ server.^73^ Accordingly, E518 was protonated in all SMO systems. For SANT1-SMO, owing to the lack of the CRD in the SANT1-SMO complex (PDB ID: 4N4W^35^), we sought to use the inactive orientation of 5L7D (inactive SMO, CRD present) instead. The SANT1-bound crystal structure (4N4W) was aligned to inac-SMO 5L7D (to maintain the same binding pose for SANT1), and the 5L7D-SANT1 starting point was used for simulations. The terminal residues were capped using neutral terminal caps Acetyl (ACE) for N-terminus and MethylAmide (NME) for the C-terminus. The proteins were embedded in a membrane bilayer using CHARMM-GUI. ^74,75^ The atomic interactions were characterized using the CHARMM36 force field. ^76,77^ The choice of CHARMM36 force field was based on studies that use CHARMM36 to simulate various G-Protein Coupled Receptors, specifically at the time of system setup.^78–81^ Use of CHARMM36m force field made noted no significant difference to the overall observations (Fig. S1). The force field parameters for synthetic ligands SAG and SANT1 were generated using ParamChem, ^82^ an automated version of CGenFF.^83,84^ Owing to presence of penalties greater than 10 assigned by CGenFF for various angles and dihedrals for both SAG and SANT1, optimization using the MP2/6-31G* QM calculations was performed. The python-based library Psi4 was used for this purpose. ^85^ Input files were generated using the web-based input generator CHARMM-GUI. ^86^ The composition of the membrane bilayer was based on lipid composition of the mice brain cerebellum^87^ - (75% POPC, 21% Cholesterol, 4% Sphingomyelin) (Table S4), to mimic physiological cerebellar membrane composition. The system was solvated using TIP3P water^88^ and 150 mM NaCl, to mimic physiological conditions. Overall the system sizes for inac-SMO, act-SMO, SAG-SMO and SANT1-SMO were 106,415, 105,971, 105,100 and 105,582 atoms with box sizes 86×86×153 Å^3^, 86×86×152 Å^3^ 86×86×152 Å^3^ and 85×85×153 Å^3^ respectively. The mass of non-protein hydrogens was repartitioned to 3.024 Da,^89^ to enable simulations with a long timestep (4 fs). Parmed, a part of the AmberTools19 package, was used for this purpose.^90^

#### Pre-Production MD

The systems were minimized for 1000 steps, using the steepest descent method, followed by minimization using the SHAKE algorithm^91^ for 14000 steps. Systems were then heated from 0-310 K using NVT conditions for 5 ns, constraining the backbone using a force constant of 10 kcal mol^−1^ Å^−2^. Systems were then equilibrated using the NPT conditions for 5ns, at 310 K and 1 bar, using similar backbone restraints. This was followed by an equilibration of 40 ns, without constraints. Apo-SMO and SANT-SMO simulations were performed using the AMBER18^90,92–95^ biomolecular simulation package. SAG-SMO simulations were performed using NAMD 2.14.^96,97^ NAMD was used in this case to aid the simulation of lone pairs associated with the Chlorine atom in SAG.

#### Production MD

Post equilibration, the GPU-accelerated pmemd.cuda package from AMBER18^90,95^ was used for production MD. Integrator timestep was set to 4fs. Periodic boundary conditions were used, and the temperature was maintained using the Langevin Thermostat. ^98^ Pressure of each of the systems was set 1 bar, and was maintained using the Monte Carlo Barostat. Particle Mesh Ewald^99^ (PME) method was used for computing long-range electrostatic interactions. SHAKE^91^ algorithm was used to restrain the Hydrogen bonds. Cutoff for nonbonded interactions was set to 10 Å. Frames were saved every 25000 steps, giving a frame rate of 100 ps between each frame. Simulations were performed using the Blue Waters super-computer(NVIDIA Tesla K20X GPUs) or our in-house computing cluster(NVIDIA GeForce GTX 980 GPUs). Apo-SMO, SAG-SMO and SANT1-SMO were simulated for a total of ~ 250*μ*s, ~36*μ*s and ~ 42*μ*s respectively.

### Adaptive sampling, feature selection and clustering

Simulating biological systems using traditional long-MD simulations to observe submillisecond dynamics is unfeasible, hence we resorted to using a parallel approach to accelerate conformational sampling, called Adaptive sampling. The simulation data after every round of simulations was clustered (feature selection explained below), and the least populated clusters were used to seed simulations for the next round. Overall, for Apo-SMO, 7 rounds of simulations were performed, collecting 30-50*μ*s per round. For SAG/SANT1-SMO, the data was collected in a similar fashion, for 3 rounds each, around 10-20 *μ*s per round. The bias introduced in the system due to selectively starting simulations from least populated clusters was eliminated by constructing a Markov state model, that estimated the reverse transition probabilities from each microstate.

The progress of the transition from inactive to active was monitored by calculating features, each of which was selected based on maximum magnitude of Δ RRCS (RRCS - Residue Residue Contact Score). RRCS is a order-parameter identifying technique that uses a flat-linear-flat scoring scheme to assign a score to contact between every residue-pair in the system.^6^ Contacts that had |Δ*RRCS*| < 3.5 (58 such distances total) (Table S5) were used. K-means clustering was used to cluster the simulation data, based on these calculated features. Clustering was performed using the pyEMMA python library. ^100^

### Markov state model construction

The high dimensionality of the data was first reduced using tICA. The tICA lagtime was optimized by observing the plateauing of the implied timescales (−2/lnλ, λ being the largest eigenvalue of the first tICA eigenvector) vs the lag time, and was set to 30 ns for the 3 systems (Apo-SMO, SAG-SMO, SANT1-SMO). The tICA reduced-dimension data was then clustered using k-means clustering. The optimum number of clusters and no. of tICA components to be used was optimized by maximizing the VAMP2 score (sum of the squares of the highest eigenvalues of the transition matrix) for a particular the number of clusters, and the convergence of the implied timescales vs the MSM lag time. (Fig. S2,S3,S4). Accordingly, the number of clusters was set to 200 (Apo-SMO) and 100 (SANT1-SMO and SAG-SMO). The MSM lag time was set to 30 ns for the three systems (Apo-SMO, SAG-SMO, SANT1-SMO). The Chapman-Kolmogorov test, which tests the validity of the MSM on 5 macrostates, was performed using pyEMMA (Fig. S5,S6,S7).

### Trajectory Analysis and Visualization

cpptraj^101^ was used for trajectory processing. VMD^102,103^ and open-source PyMOL^104^ were used to visualize and render images. MDTraj^105^ was used for computing all order parameters. All plots were made using matplotlib ^106^ and seaborn^107^ python libraries. Numpy^108^ was used for numerical computations. The salt-bridge based contacts were discovered by extracting probability-weighted 10000 frames from clusters the in Inactive, I^1–3^ and Active states, using cluster probabilities from the MSM. They were analyzed for unique contacts using GetContacts. ^109^ Tunnel radii for analysis of effect of SAG and SANT1 were calculated using HOLE. ^67^

### Mutual Information Calculations

Mutual Information for describing the allosteric pathways was computed using mdentropy,^110^ using the DihedralMutualInformation function. Analysis was performed on 10000 frames each extracted from Apo-SMO, SANT1-SMO and SAG-SMO data. The frames were chosen based on the predicted MSM probabilities, to represent the entire ensemble. A graph was constructed from the computed Mutual Information, and residues with C-*α* distances < 10 Å were considered to be connected by an edge. The weight of each edge was assigned as MI = MI_*max*_ - MI_*ab*_, with the MI_*max*_ as the maximum mutual information computed among two residues in a protein, and MI_*ab*_ was the mutual information computed between residue pair ab. Edges with MI < MI_*avg*_ were not considered. This methodolody thus adapted has been discussed previously.^69–71^ The caveats and limitations presented by the methodology – the presence of global dynamics independent of the local dynamics being explored by the limited simulation data^111^ have been resolved by using long timescale simulations. Allosteric pathways were computed by calculating the shortest paths between 2 nodes, (in our case E518 and W339) using Dijkstra’s algorithm.^112^ NetworkX,^113^ a python library was used for graph-construction, visualization and computing shortest paths.

## Supporting information

Supplementary text, figures, methods and references

## Data Availability

Stripped trajectories and corresponding parameter files have been uploaded to [Box](https://uofi.box.com/s/c12lwov1mtrncm30hydu9iqhb45dcwx4). Scripts used for MSM construction and trajectory analysis have been uploaded to [github](https://github.com/ShuklaGroup/Bansal_et_al_Smoothened_Activation_2022).

## Author Contributions

D.S. designed the research. P.D.B. performed simulations. P.D.B and S.D. analyzed the data. P.D.B. wrote the manuscript with inputs from S.D. and D.S.

## Acknowledgments

The authors thank The Blue Waters Petascale Computing Facility and National Center for Supercomputing Applications, which is supported by the National Science Foundation (awards OCI-0725070 and ACI-1238993) and the state of Illinois. Blue Waters is a joint effort of the University of Illinois at Urbana-Champaign and its National Center for Supercomputing Applications. D.S. acknowledges support from NIH grant R35GM142745 and Cancer Center at Illinois for their support. P.D.B thanks Austin Weigle and Jiming Chen of the Shukla Group at University of Illinois for the valuable insights throughout the course of this study.

