## Supplementary text, figures, methods and references for "Activation mechanism of the human Smoothened receptor"

### List of Figures


S1 CHARMM36mfigure . . . . . . . . . . . . . . . . . . . . . . . . . . . S-4
S2 MSM construction for Apo-SMO . . . . . . . . . . . . . . . . . . . . S-5
S3 MSM construction for SANT1-SMO . . . . . . . . . . . . . . . . . . . S-6
S4 MSM construction for SAG-SMO . . . . . . . . . . . . . . . . . . . . S-7
S5 MSM validation for Apo-SMO . . . . . . . . . . . . . . . . . . . . . S-8
S6 MSM validation for SANT1-SMO . . . . . . . . . . . . . . . . . . . . S-9
S7 MSM validation for SAG-SMO . . . . . . . . . . . . . . . . . . . . . S-10
S8 Roundwise data collection for adaptive sampling in Apo-SMO . . . . . . . . S-11
S9 Apo-SMO tICA plots outlining the slowest processes . . . . . . . . . . . . S-12
S10 TICA plots for SANT1-SMO and SAG-SMO . . . . . . . . . . . . . . . . S-13
S11 TICA Scatterplots show residue-correlated movements with activation on tICA space . . . . . . . . . . . . . . . . . . . . . . . . . . . . . . S-14
S12 Multiple Sequence Alignment of Class F receptor Transmembrane Domains . S-15
S13 Error in the free-energy plots . . . . . . . . . . . . . . . . . . . . . S-16
S14 Qualitative comparison in the structural commonalities between Class A and Class F GPCRs . . . . . . . . . . . . . . . . . . . . . . . . . . . S-17
S15 Difference in the polarity of residues in the CRD . . . . . . . . . . . . . S-18
S16 Rearrangement of the salt-bridges on the CRD-TMD interface. Inset shows the particular salt bridges involved in SMO activation. . . . . . . . . . . S-19
S17 Multiple Sequence Alignment for the Class F CRD . . . . . . . . . . . . S-20
S18 Rotation of CRD on activation . . . . . . . . . . . . . . . . . . . . . S-21
S19 Hydrophobic tunnel inside SMO . . . . . . . . . . . . . . . . . . . . S-22
S20 Pathway to the CRD . . . . . . . . . . . . . . . . . . . . . . . . . S-22
S21 Lateral movement of SANT1 . . . . . . . . . . . . . . . . . . . . . S-23
S22 Cholesterol Distributions . . . . . . . . . . . . . . . . . . . . . . . S-24
S23 Tunnel Opening between TM2 and TM3 . . . . . . . . . . . . . . . . . S-25

S24 Tunnel Opening between TM2 and TM3 . . . . . . . . . . . . . . . . . . . . S-26
S25 Helical displacement of TM6 by SANT1 . . . . . . . . . . . . . . . . . . . . S-27

### List of Tables

S1 Round wise data collection for Apo-SMO . . . . . . . . . . . . . . . . . . . . S-28
S2 Round wise data collection for SAG-SMO and SANT1-SMO . . . . . . . . . . S-28
S3 Modelled residues in 5L7D-inactive-Apo-SMO . . . . . . . . . . . . . . . . . S-29
S4 Lipid Composition . . . . . . . . . . . . . . . . . . . . . . . . . . . . . S-29
S5 Adaptive Sampling metrics . . . . . . . . . . . . . . . . . . . . . . . . . S-30

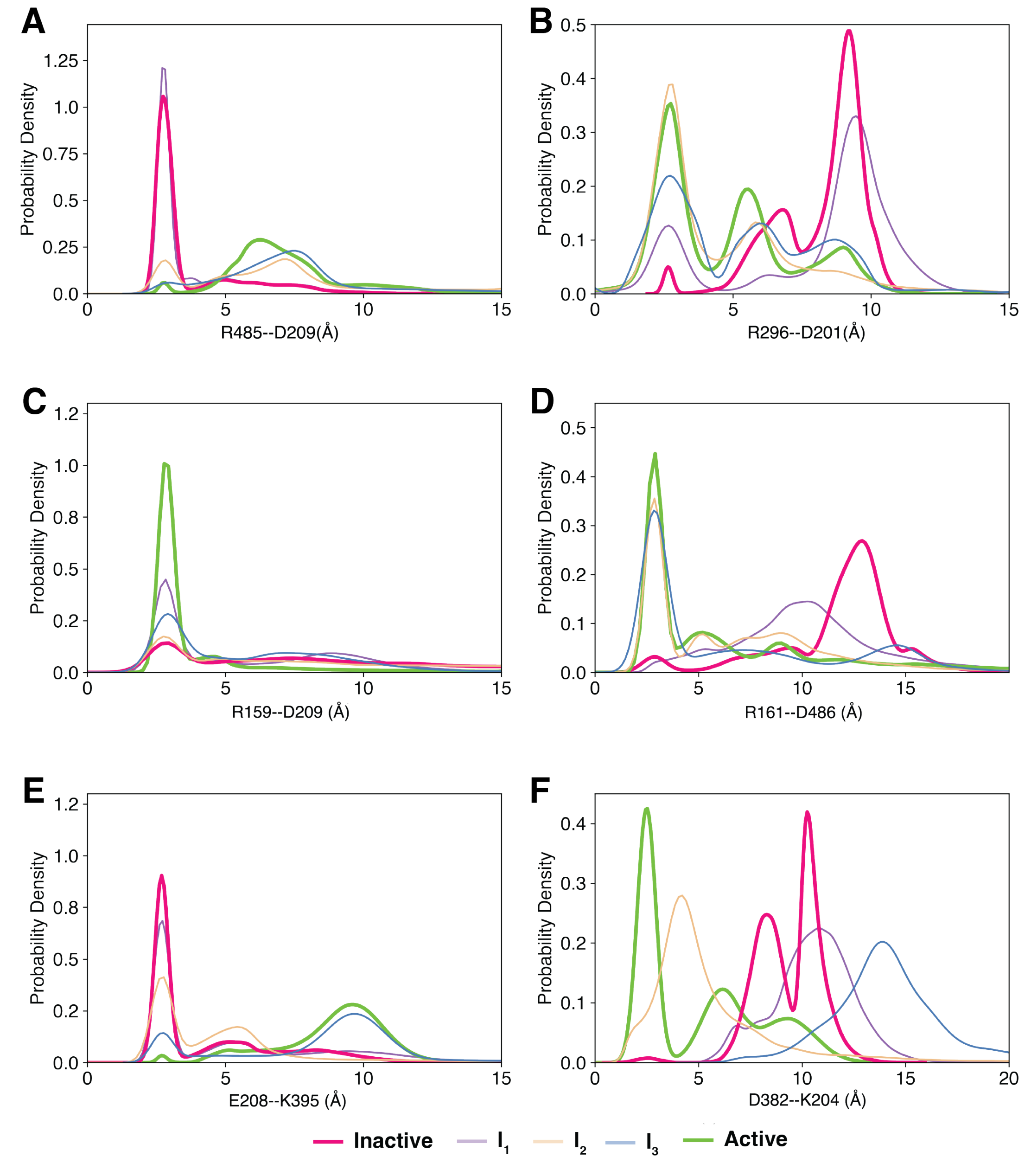


Fig. S1: Figure 3 reconstructed using CHARMM36m force field. Use of CHARMM36m force field made noted no significant difference to the overall observations made using CHARMM36 force field.

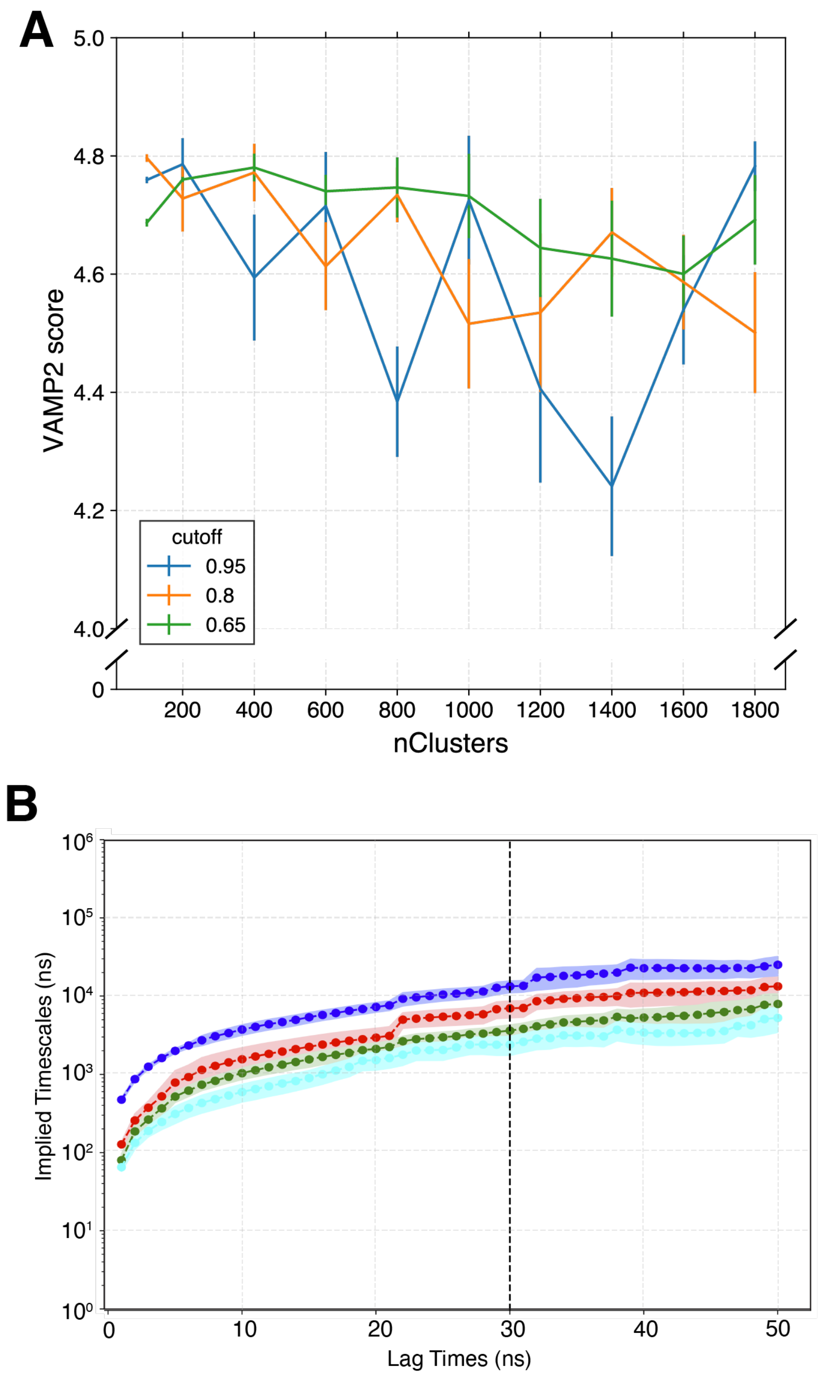


Fig. S2: MSM construction for Apo-SMO. (A) VAMP2 score v/s nClusters used to cluster the data, as a function of varying variational cutoffs for choosing the number of tICA (time Independent Component Analysis) components. For final MSM construction, 200 clusters with a 0.95 variational cutoff (corresponding to 42 tICA components) was chosen to construct the MSM. (B) Implied Timescales v/s MSM lag time for the MSM with 200 clusters and 42 tICA components. A lagtime of 30 ns was chosen for construction of the final MSM.

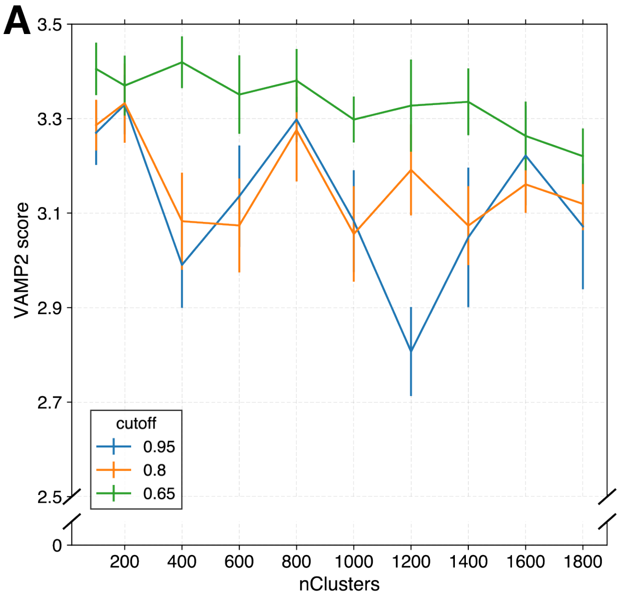


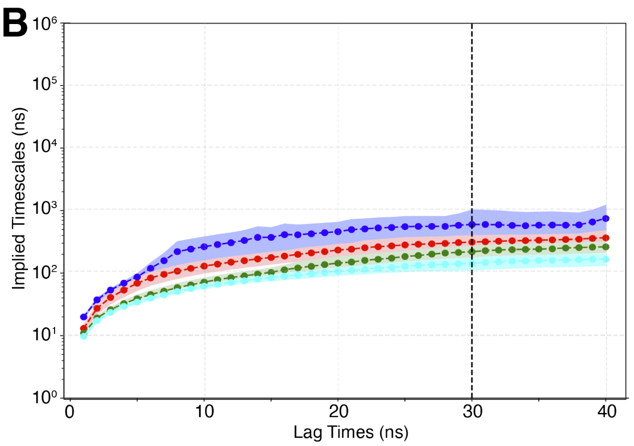


Fig. S3: MSM construction for SANT1-SMO. (A) VAMP2 score v/s nClusters used to cluster the data, as a function of varying variational cutoffs for choosing the number of tICA (time Independent Component Analysis) components. For final MSM construction, 100 clusters with a 0.95 variational cutoff (corresponding to 34 tICA components) was chosen to construct the MSM. (B) Implied Timescales v/s MSM lag time for the MSM with 100 clusters and 34 tICA components. A lagtime of 30 ns was chosen for construction of the final MSM.

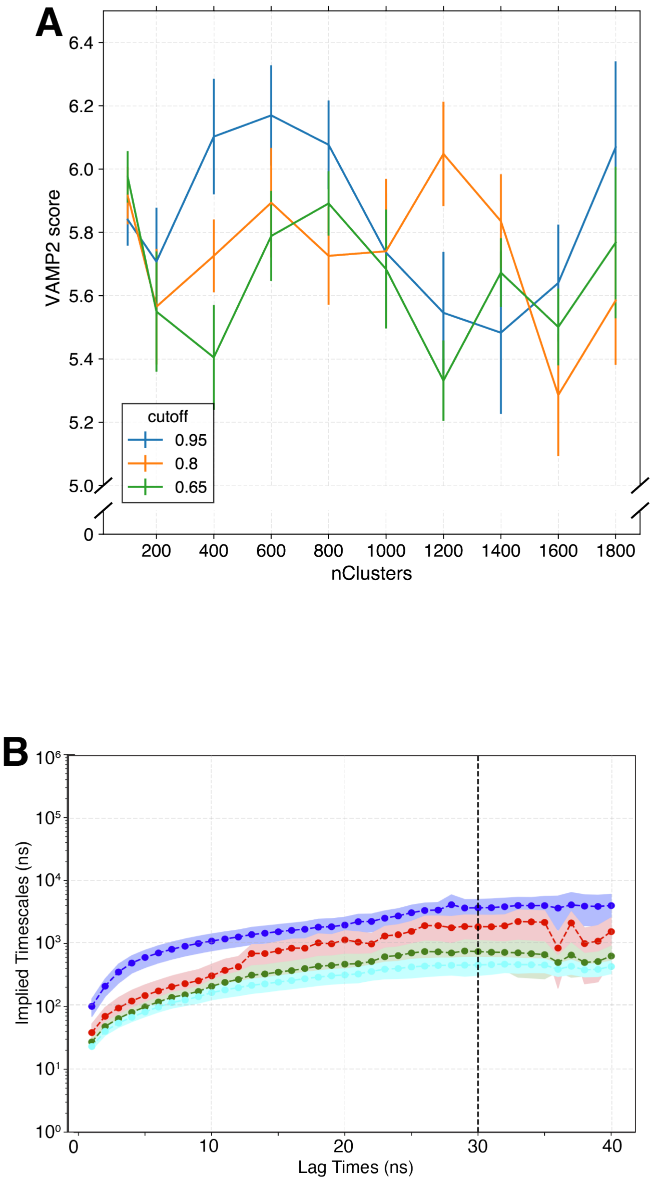


Fig. S4: MSM construction for SAG-SMO. (A) VAMP2 score v/s nClusters used to cluster the data, as a function of varying variational cutoffs for choosing the number of tICA (time Independent Component Analysis) components. For final MSM construction, 100 clusters with a 0.95 variational cutoff (corresponding to 34 tICA components) was chosen to construct the MSM. (B) Implied Timescales v/s MSM lag time for the MSM with 100 clusters and 34 tICA components. A lagtime of 30 ns was chosen for construction of the final MSM.

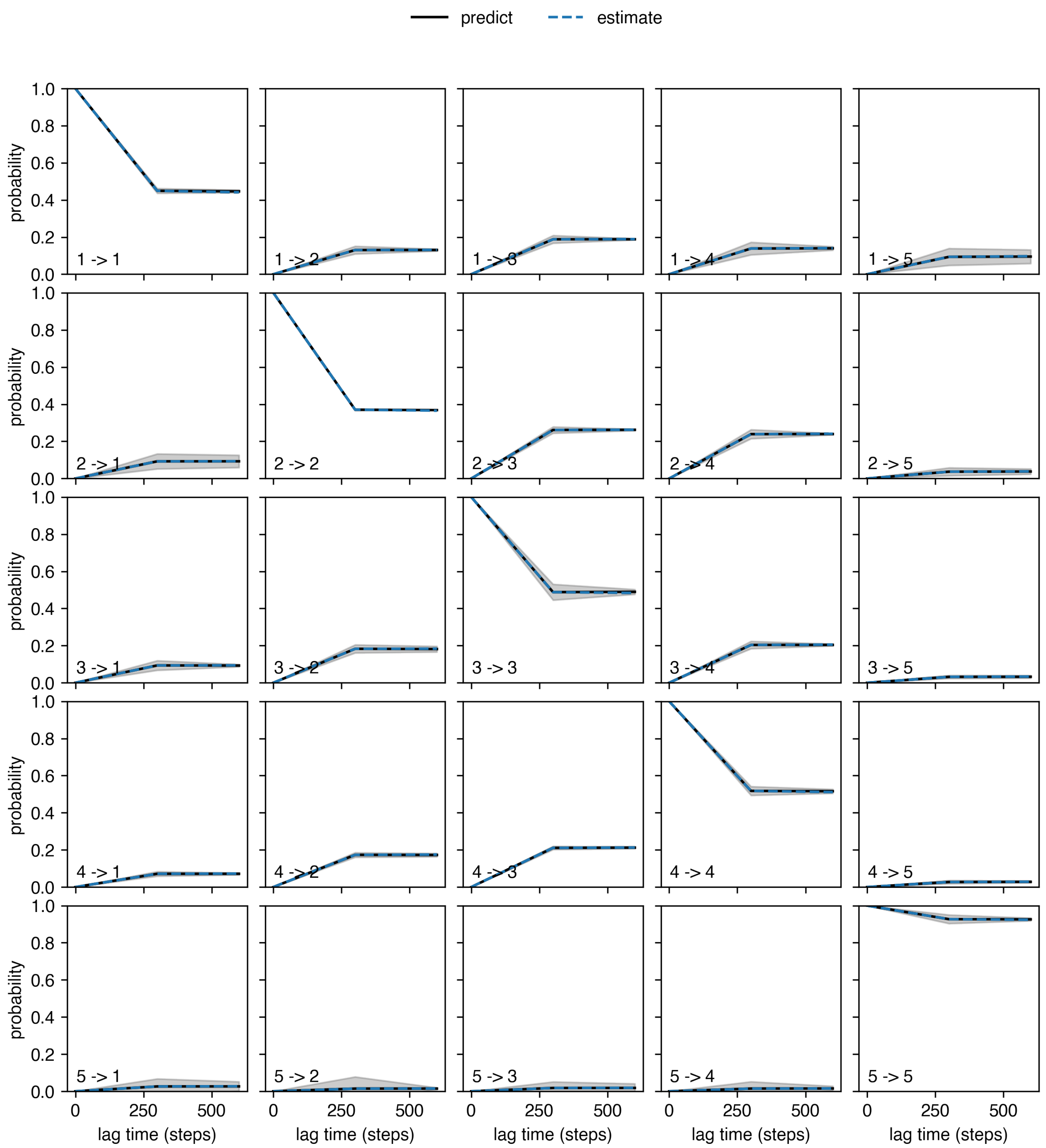


Fig. S5: MSM validation for Apo-SMO. Chapman-Kolmogorov test performed for 5 macrostates for Apo-SMO. Chapman-Kolmogorov test was implemented using the pyEMMA package.[1]

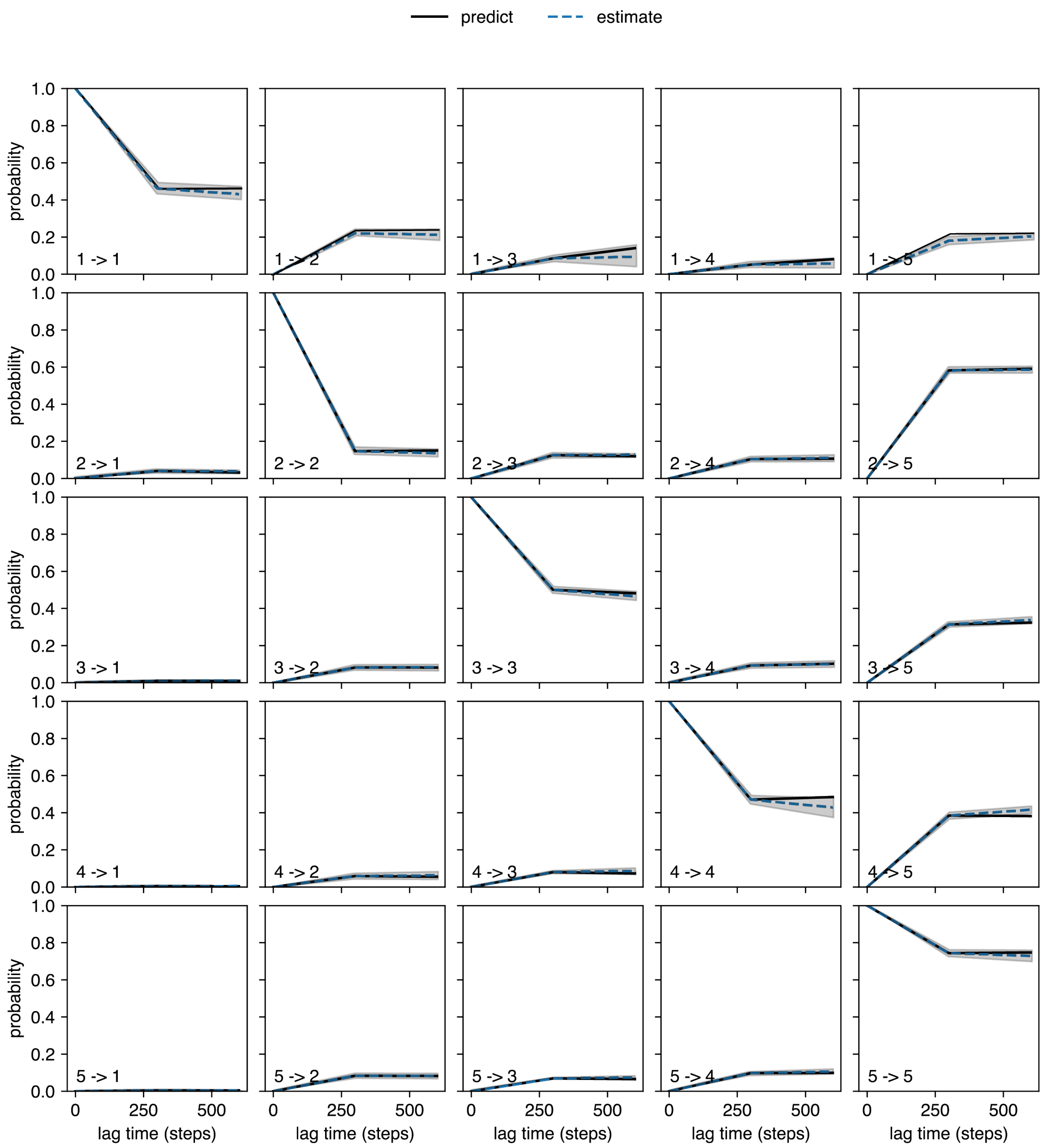


Fig. S6: MSM validation for SANT1-SMO. Chapman-Kolmogorov test performed for 5 macrostates for SANT1-SMO. Chapman-Kolmogorov test was implemented using the pyEMMA package.[1]

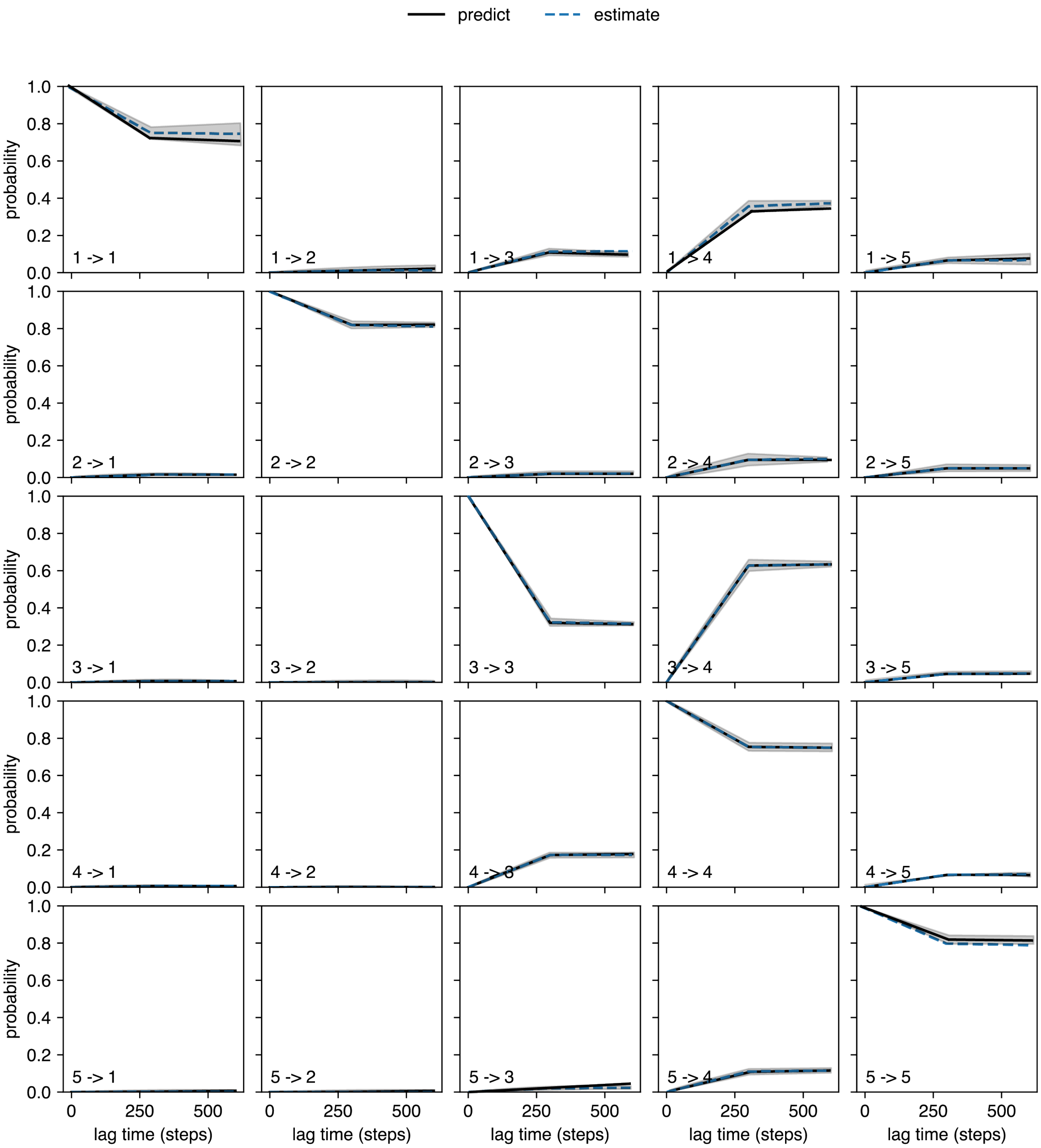


Fig. S7: MSM validation for SAG-SMO. Chapman-Kolmogorov test performed for 5 macrostates for SAG-SMO. Chapman-Kolmogorov test was implemented using the pyEMMA package.[1]

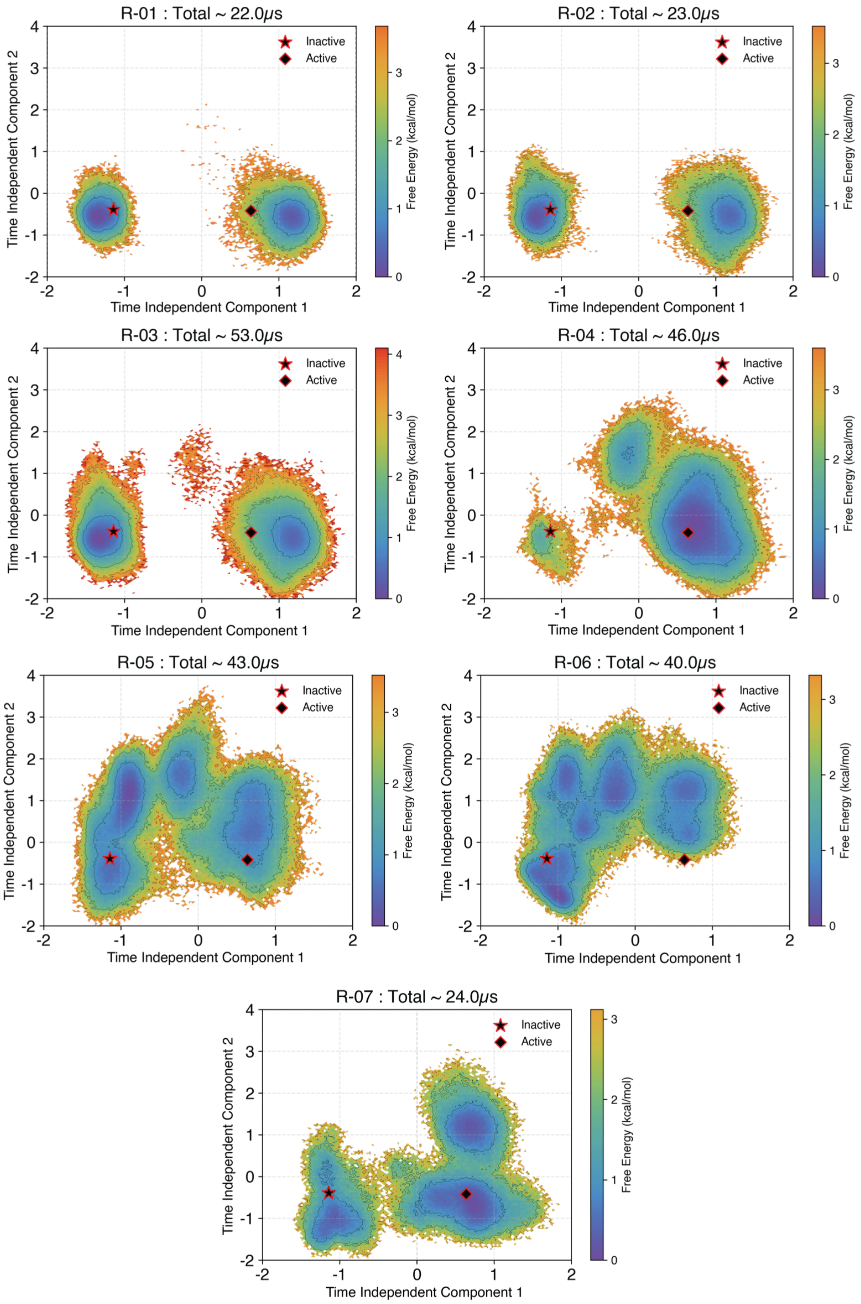


Fig. S8: Roundwise data collection for adaptive sampling in Apo-SMO. The 2 startpoints and amount of data collected in that round are mentioned on each plot. A total of 7 rounds of sampling was performed.

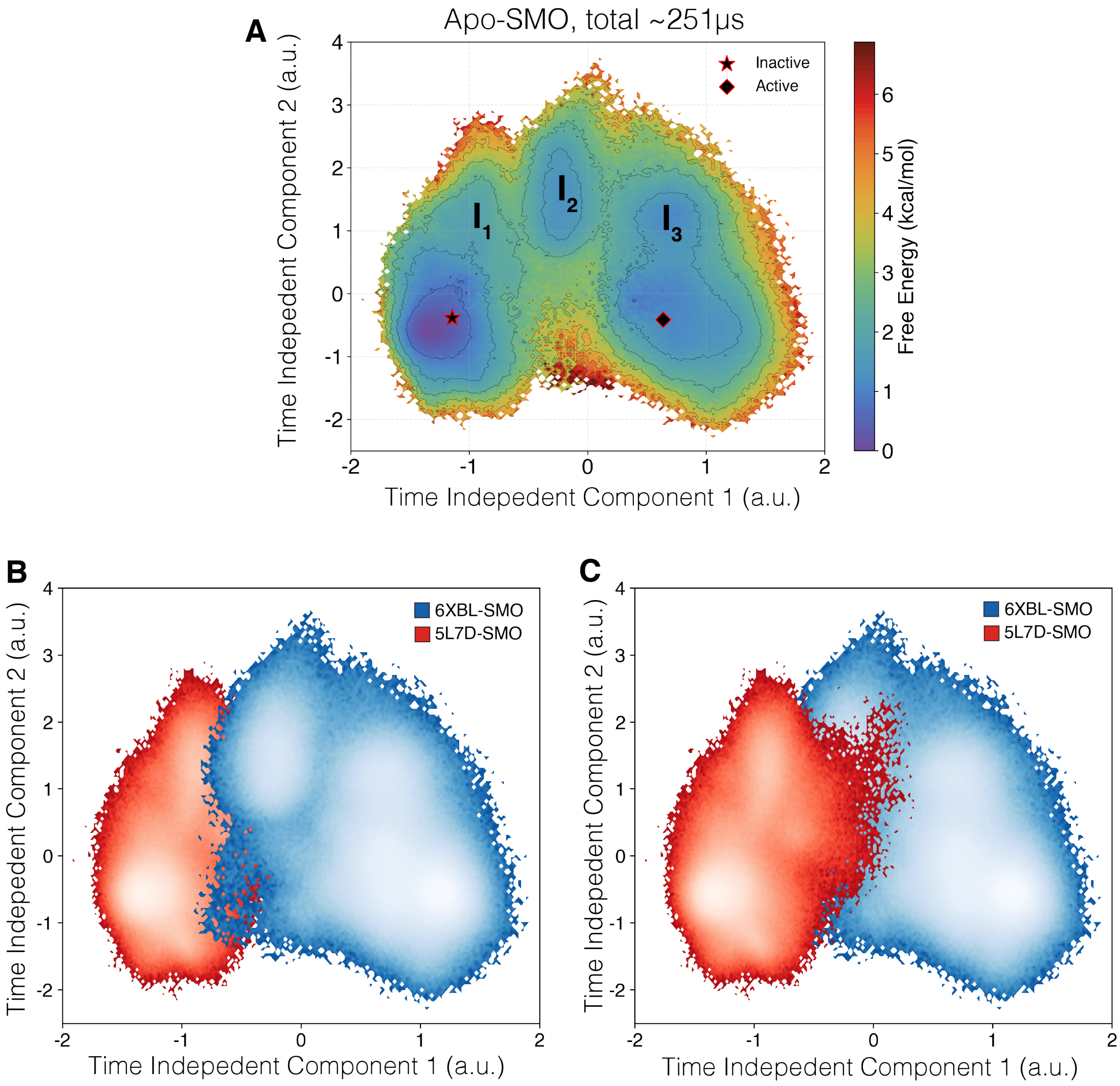


Fig. S9: Apo-SMO tICA plots outlining the slowest processes. (A) Projection of MSM weighted simulation data for Apo-SMO on the tICA space, using the 2 slowest components. The inactive and active starting points are marked. The inactive and active starting points show a free energy difference of $\sim$ 1 kcal.mol$^{-1}$. Intermediate states $I_1$, $I_2$ and $I_3$ are as marked. The intermediate states $I_{1-3}$ were defined based on metastable basins and free energy barriers associated with transitioning from an inactive to an active state. A cutoff of 1.8 kcal/mol was used to separate one basin from another. (B,C) Projection of the data for each starting point shown separately on the tICA landscape. The two islands show overlap, indicating transitions that span the slowest component. Red denotes the data collected from the inactive starting structure, while Blue denotes the data collected from the active starting structure.

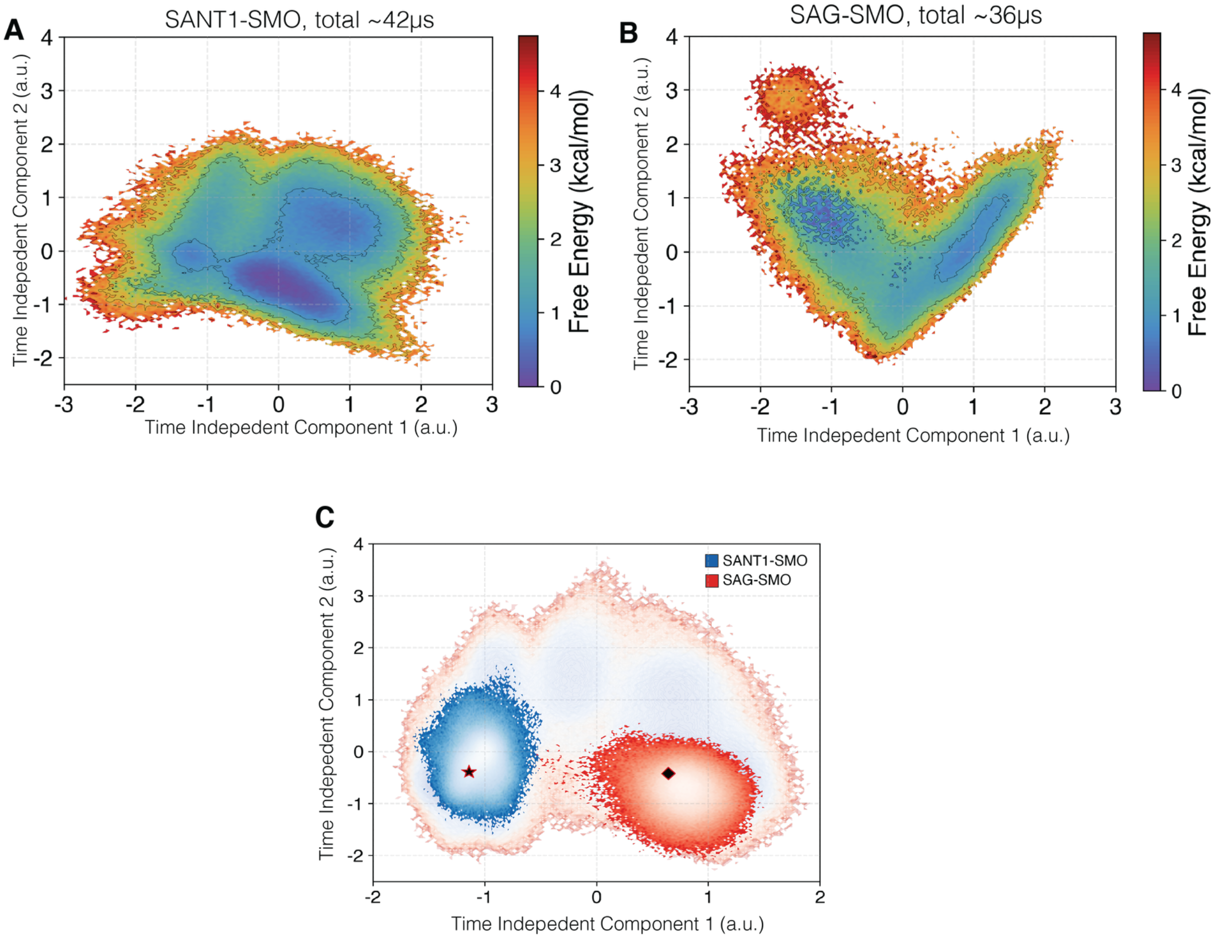


Fig. S10: Projection of MSM weighted simulation data for (A) SANT1-SMO and (B) SAG-SMO on the tICA space, using the 2 slowest components. (C) The same data as (A) and (B), when projected on the tICA space defined by Apo-SMO. The inactive and active structures are marked as star and diamond, respectively.

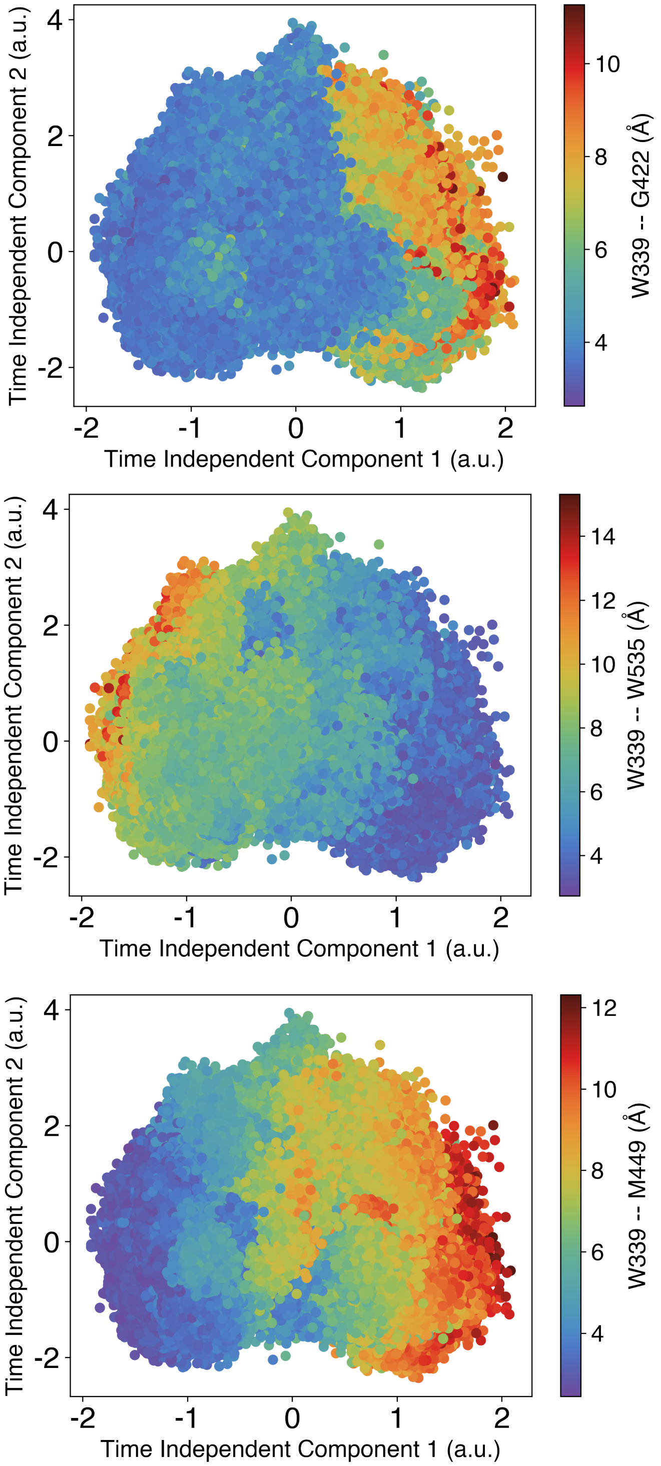


Fig. S11: TICA Scatterplots show residue-correlated movements with activation on tICA space. Scatterplots showing the correlation of the intracellular residue distances W339-G422 (TM3-TM5), W339-W535 (Ionic Lock) and W339-M449 (TM3 - TM6), projected on the tICA landscape. tIC1 corresponds to SMO activation, hence these distances are integral to SMO activation.

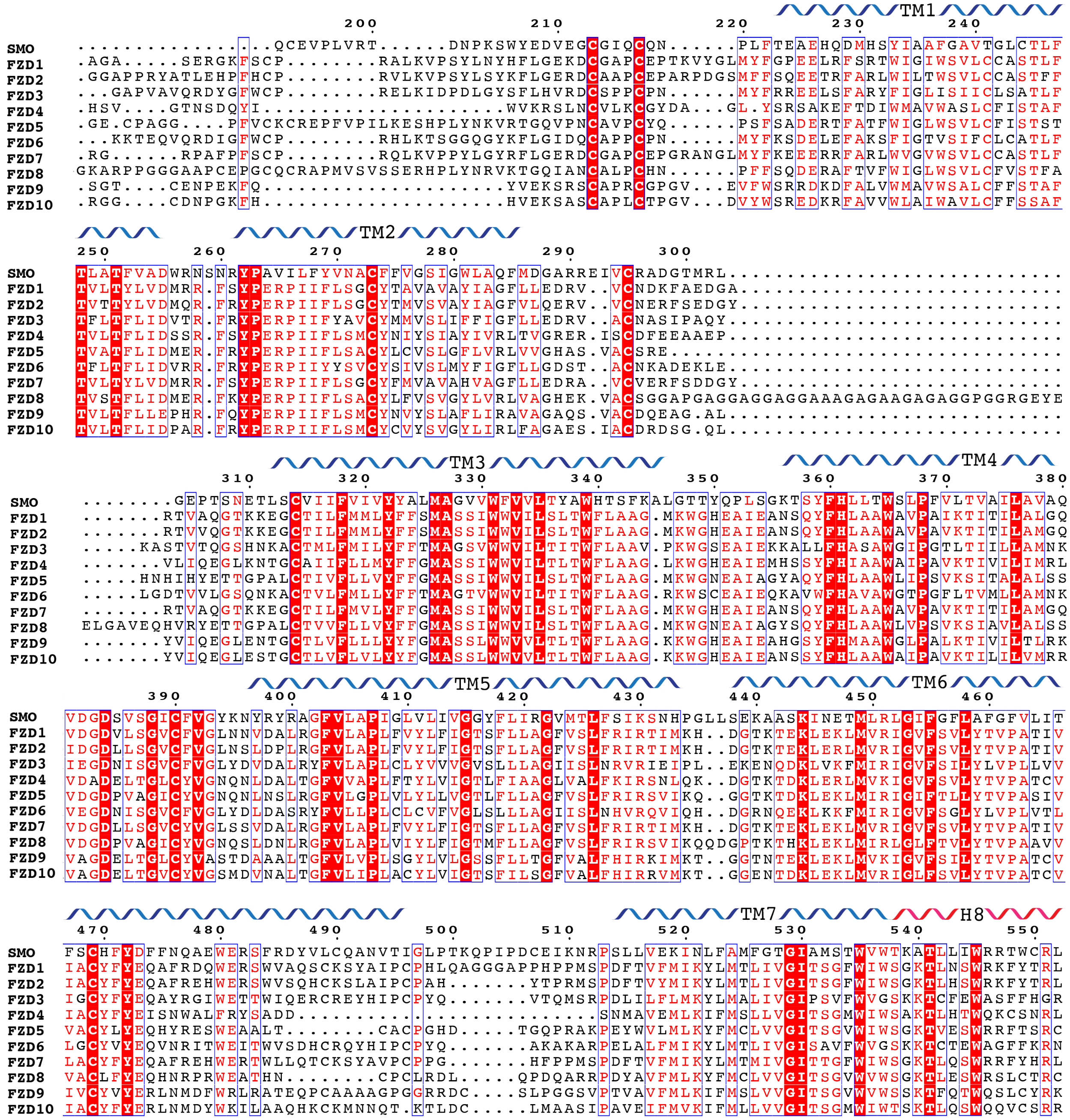


Fig. S12: Multiple Sequence Alignment of Class F receptor Transmembrane Domains. Residues highlighted in red are conserved across the entire family.

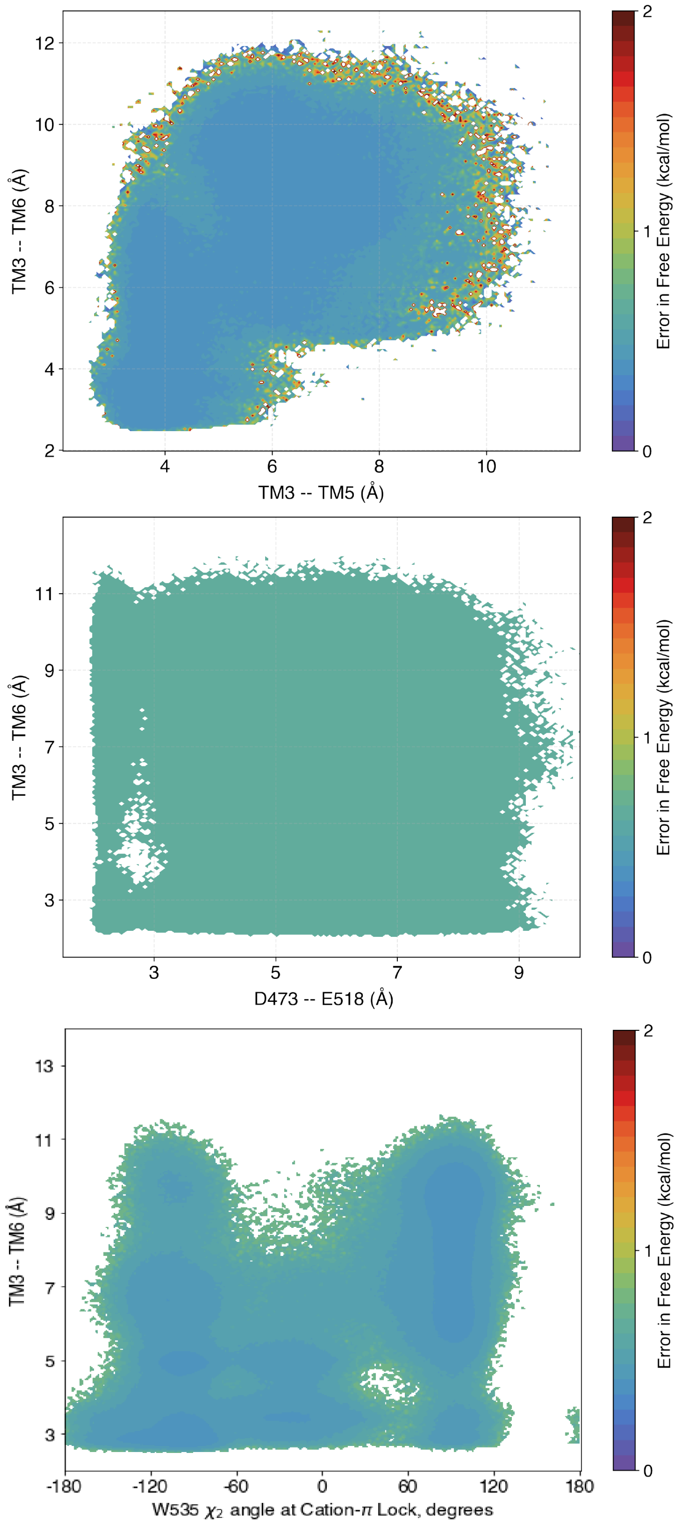


Fig. S13: Error in the free-energy plots discussed in Fig 2 of the main text. Errors calculated using bootstrapping - 80% of the data was used in 200 iterations to produce the error.

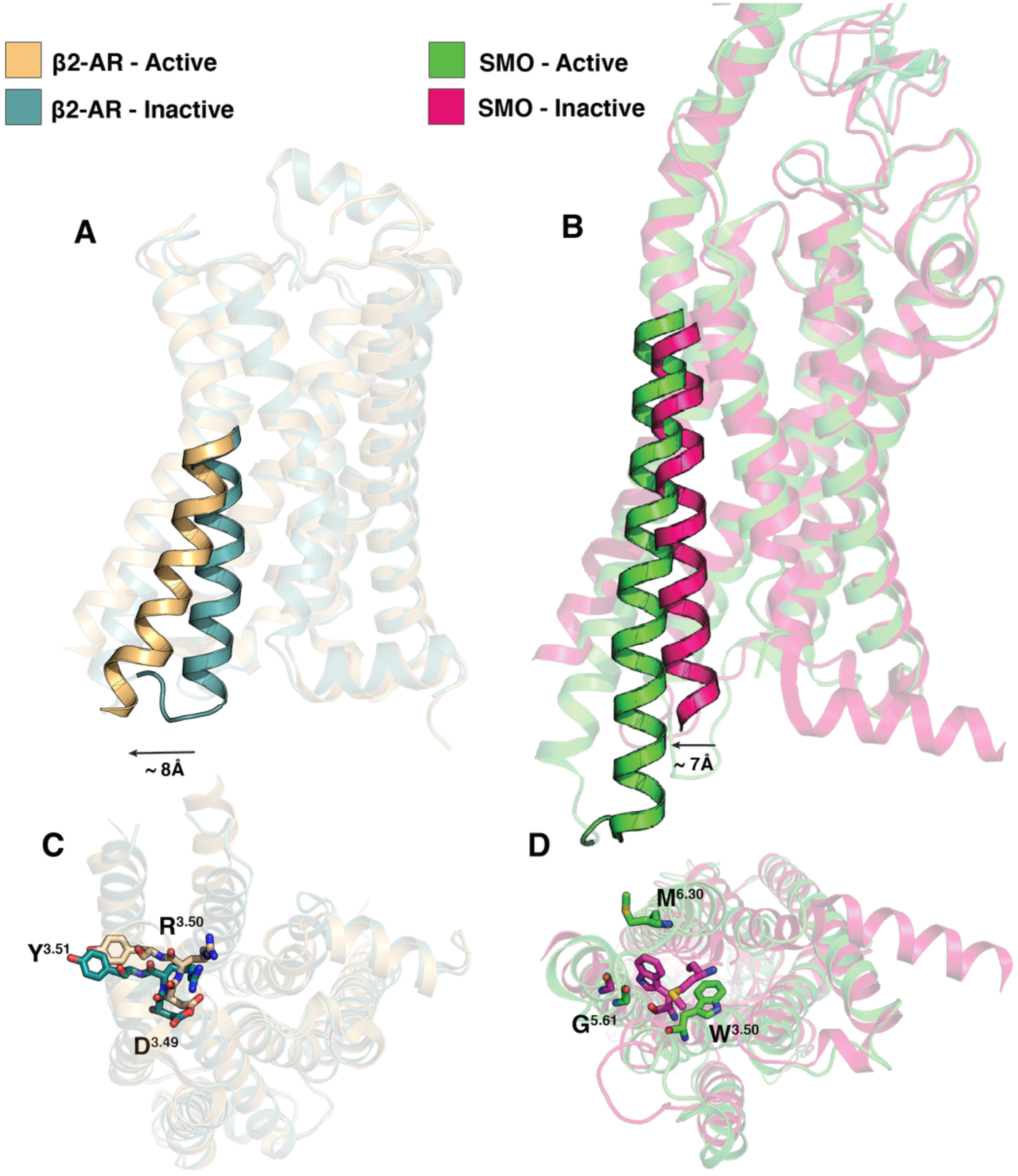


Fig. S14: Qualitative comparison in the structural commonalities between Class A and Class F GPCRs. $\beta_2$-AR (A), a class A GPCR, shows the canonical outward movement of TM6 corresponding to receptor activation.[2,3] SMO (B) also shows the corresponding outward movement. (C,D) Intracellular view of the D-R-Y (C) and W-G-M (D) motifs.

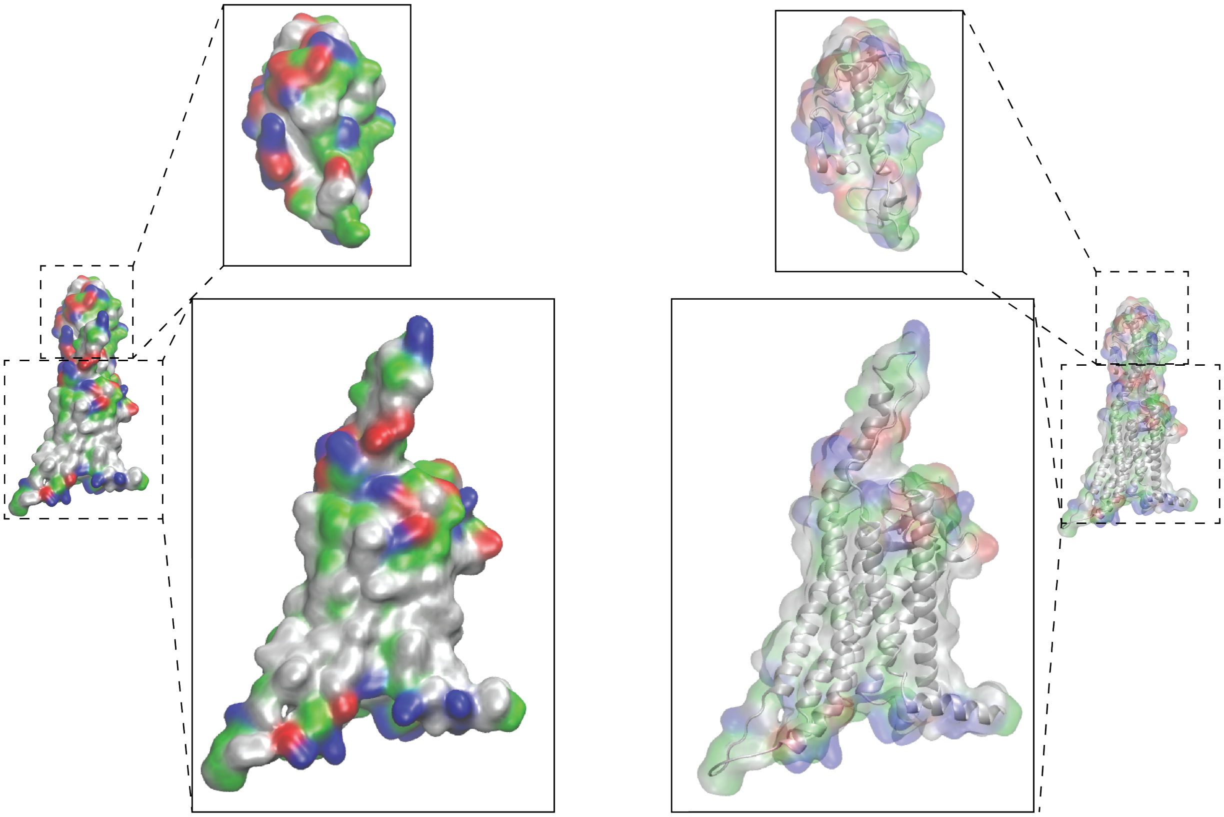

Fig. S15: Difference in the polarity of residues between the CRD (top) vs TMD (bottom). Residues colored in red-blue are polar, while green-white residues show the hydrophobic residues.

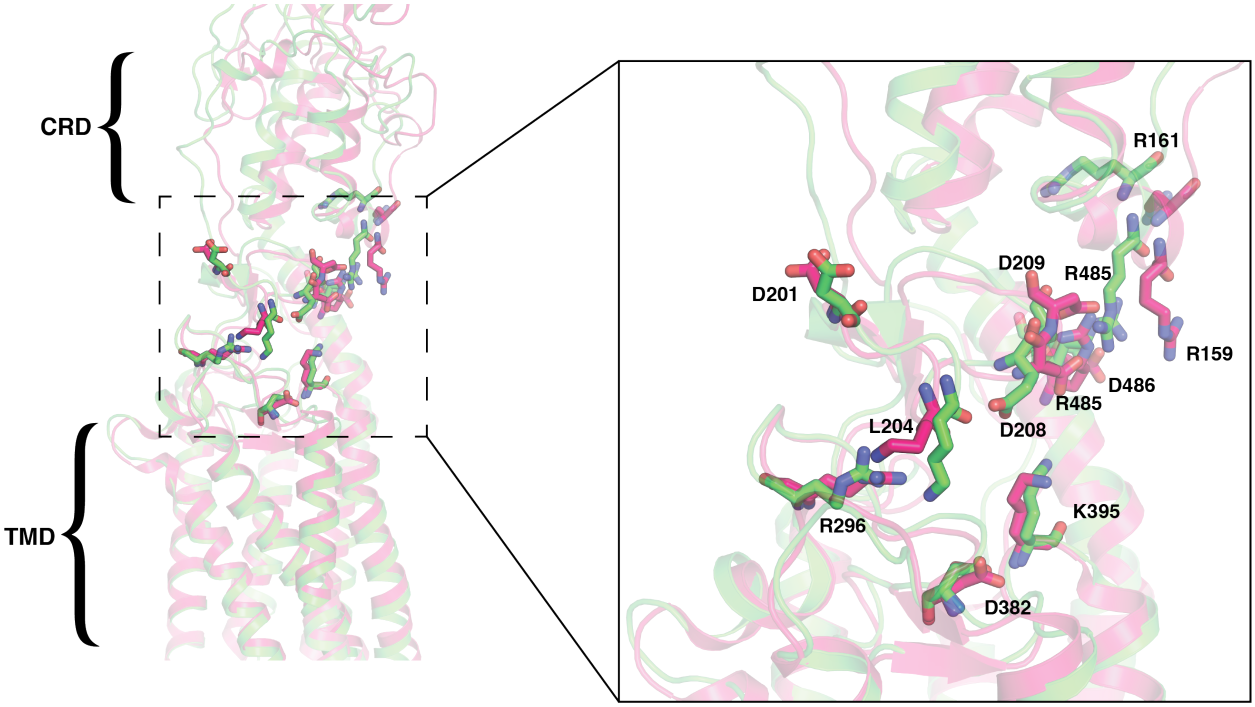


Fig. S16: Rearrangement of the salt-bridges on the CRD-TMD interface. Inset shows the particular salt bridges involved in SMO activation.

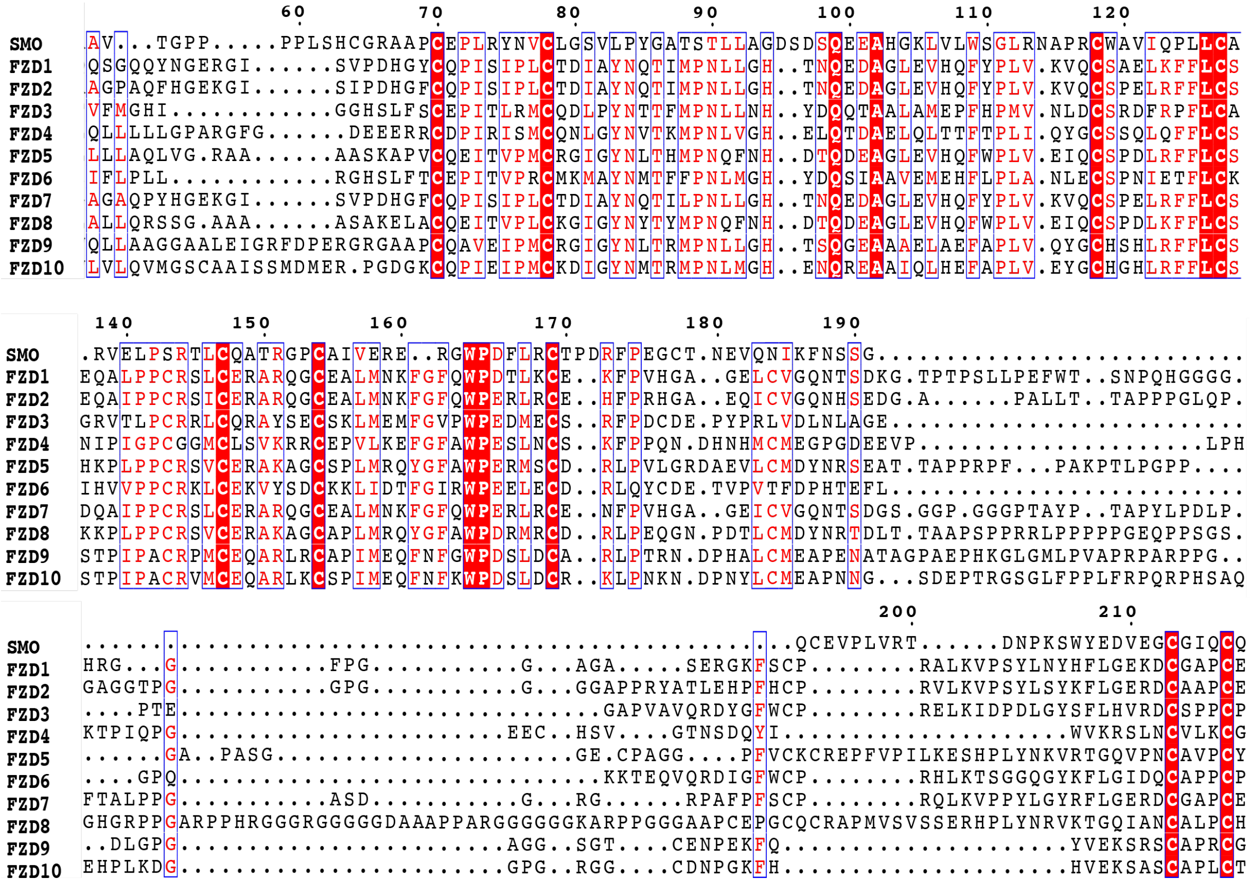

Fig. S17: Multiple Sequence Alignment for the CRD of Class F receptors. The alignment was perfomed using ESPript3 web server[4,5]

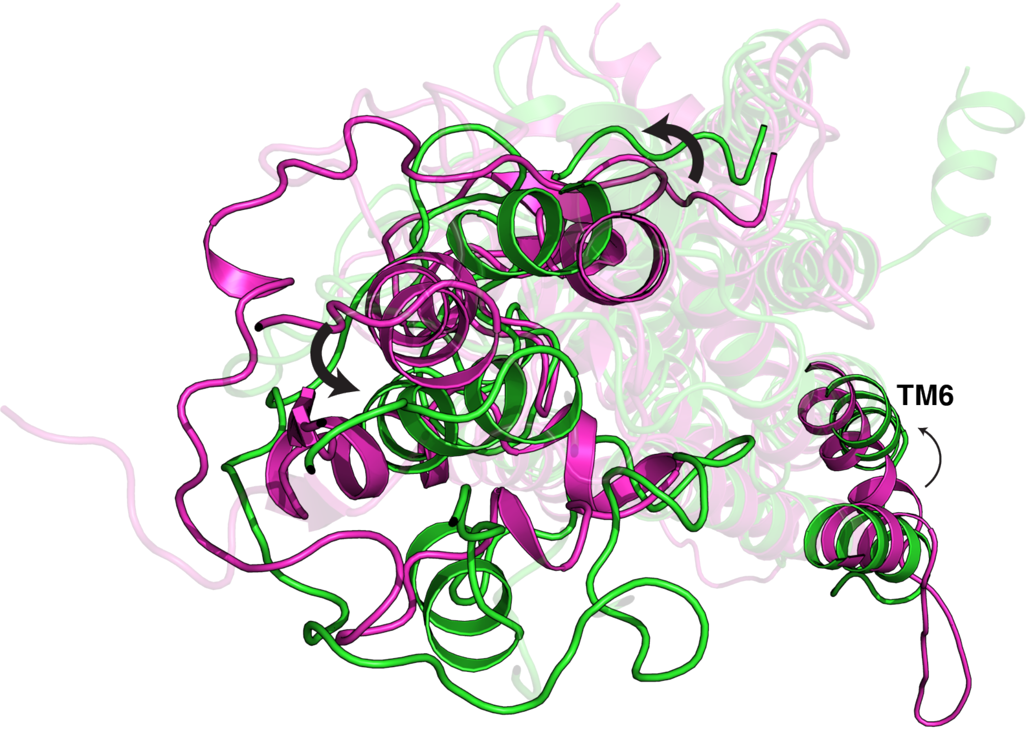


Fig. S18: Rotation of CRD on activation. (Pink-Inactive SMO; Green-Active SMO) CRD undergoes a reorientation during SMO activation, which is characterized by the outward movement of TM6.

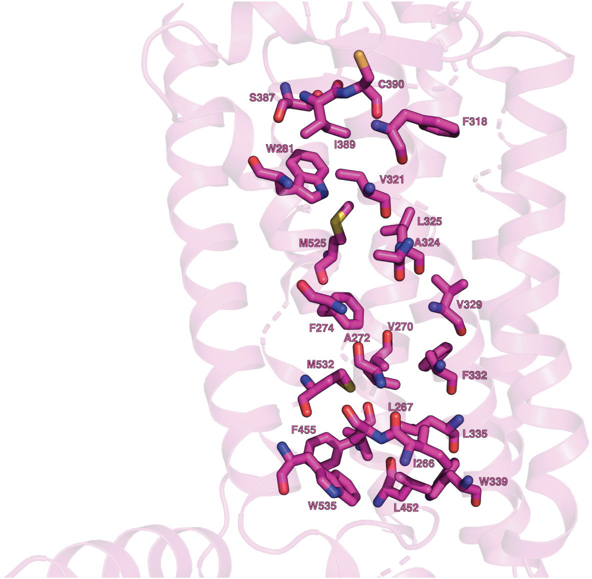


Fig. S19: Hydrophobic tunnel inside SMO. The tunnel inside SMO consists of primarily hydrophobic residues, lining the tunnel from the extracellular end (top) to the intracellular end (bottom)

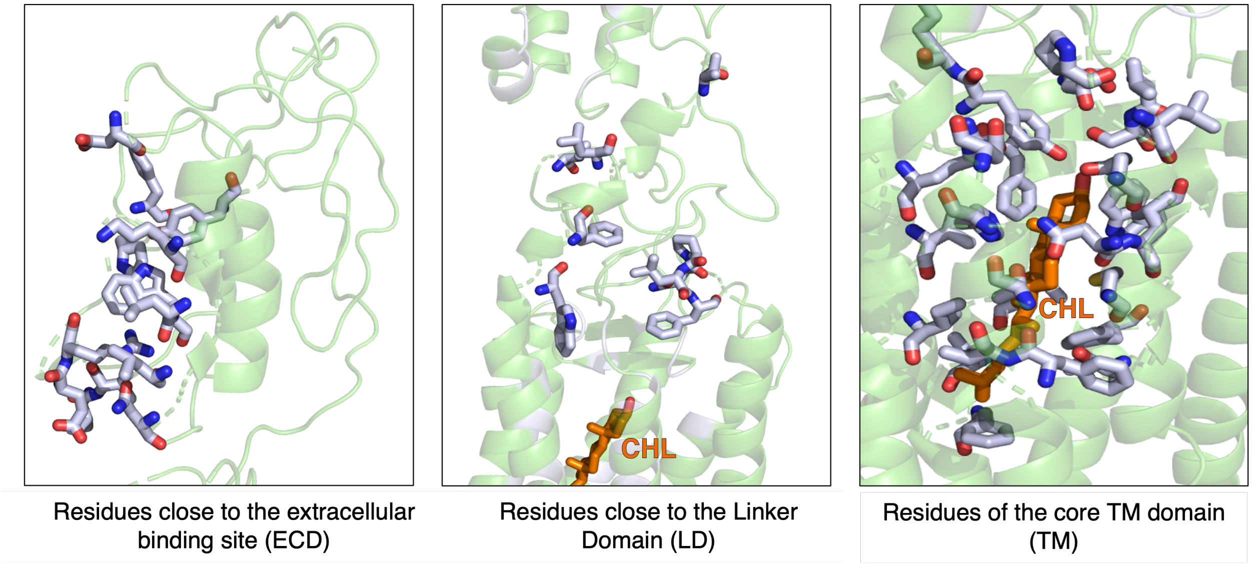


Fig. S20: Pathway to the CRD for proposed cholesterol transport. Various residues line the pathway between the entry-point of cholesterol from the membrane to the binding site, in the CRD. These consist of residues close to the extracellular binding site in the ECD, in the linker domain (LD) which connects the CRD to the TMD, and in the core TM domain. The residues were determined based on the proximity of the sterol in multiple sterol-bound resolved structures of SMO (PDB ID 6XBL, 6XBM[6] and 5L7D.[7]

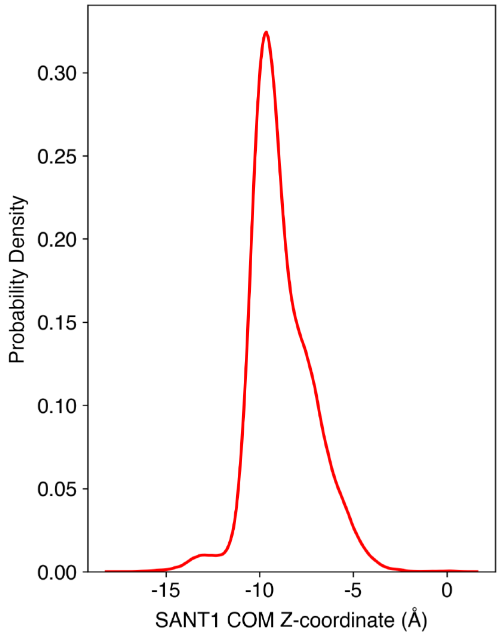


Fig. S21: Lateral movement of SANT1 center of mass. Simulations show a minimal lateral movement of SANT1 across the hydrophobic tunnel, suggesting that SANT1 operates as an antagonist by sterically blocking the tunnel.

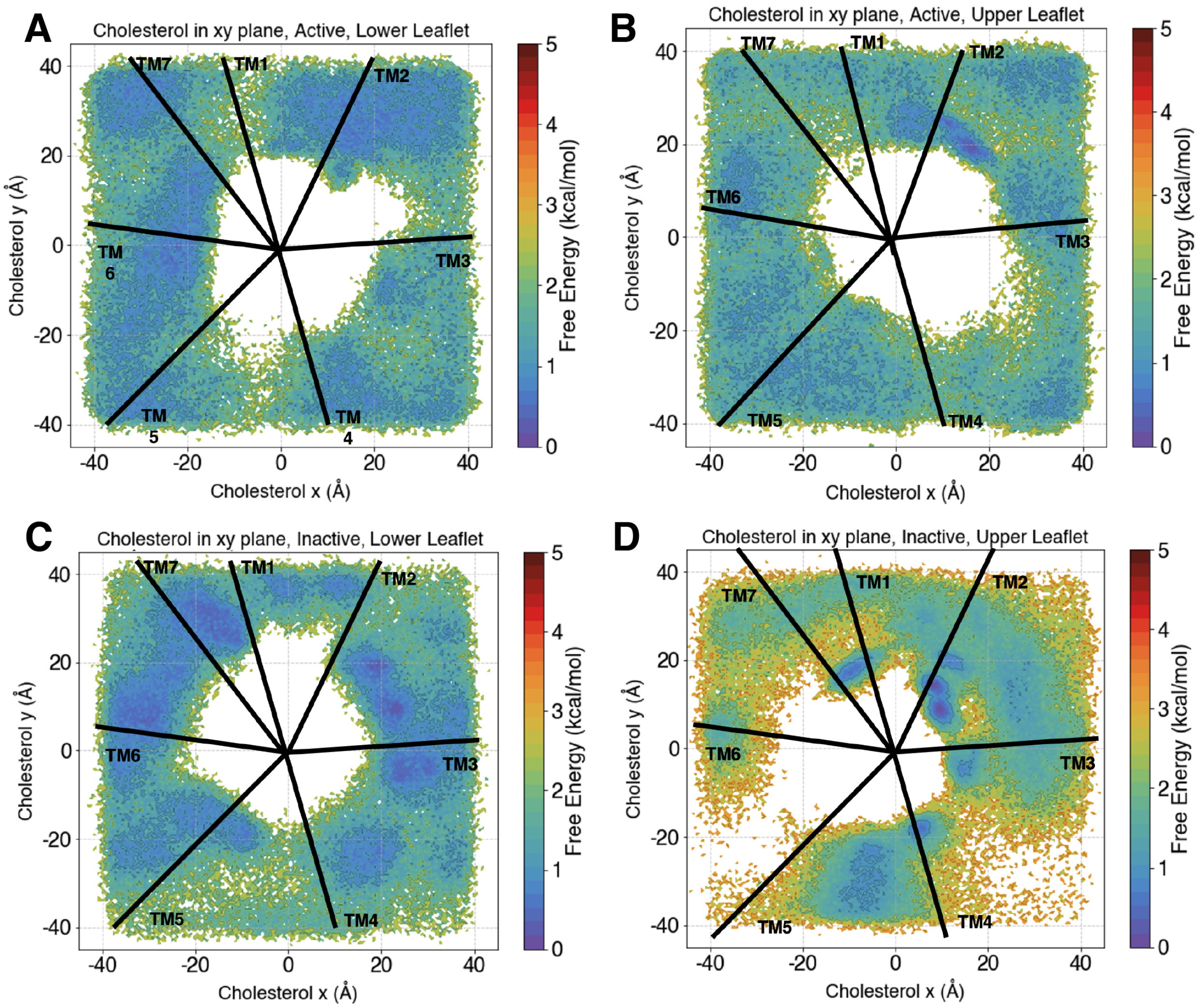


Fig. S22: Observed cholesterol densities in the membrane during Apo-SMO simulations. The plots show the distribution of the cholesterol in the membrane, with the white space in the middle occupied by SMO. (A-C) Cholesterol shows a uniform distribution in the Lower leaflet regardless of the conformation, as well as the upper leaflet in the active state. (D) Cholesterol, however, does show a propensity to cluster outside the area between TM2 and TM3 in the upper leaflet, showing a conformational dependence on the cholesterol distribution. The black lines show the average position of the helix in the membrane as seen from above, looking directly into the core of the protein.

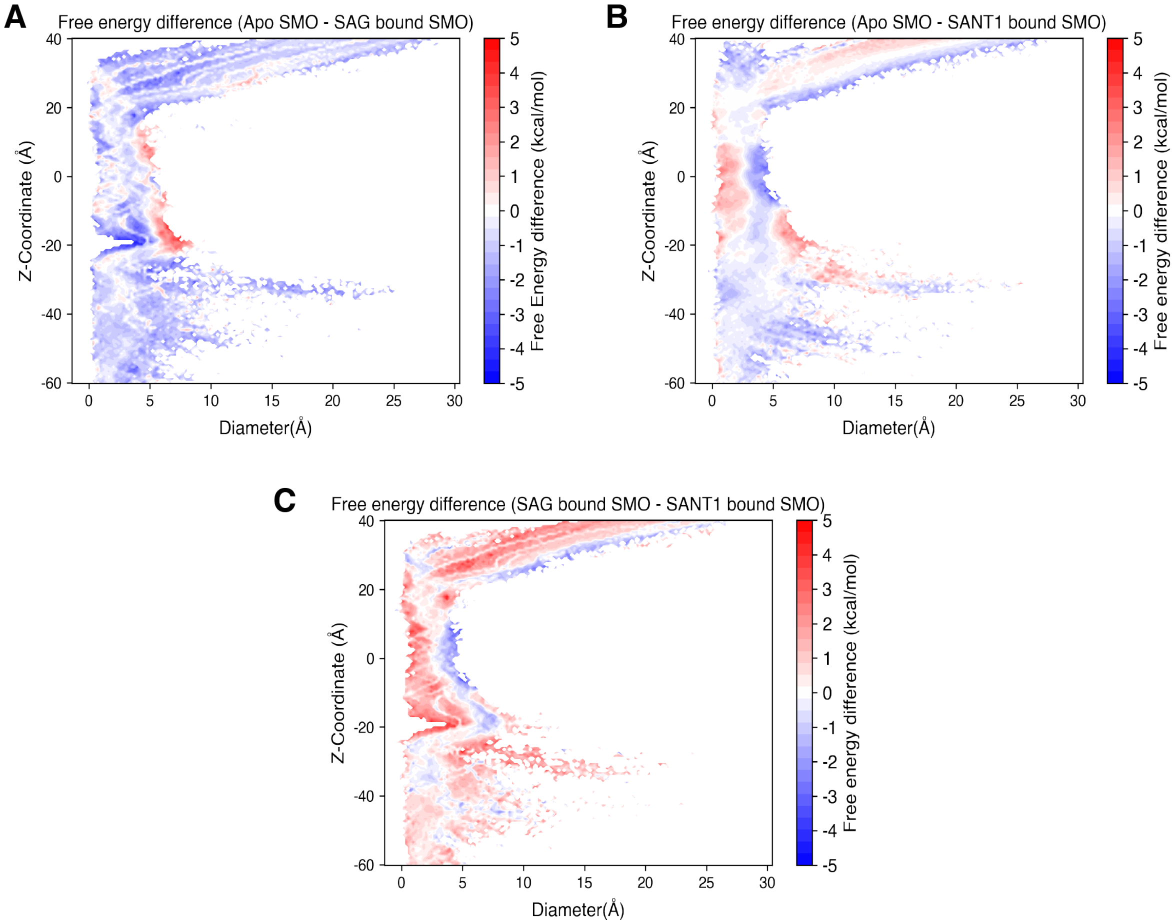


Fig. S23: (A) Probability density of Z-coordinate of tunnel opening. Simulations show that the tunnel opens in the upper leaflet of the membrane, z ∼ -22. (B) The Y v/s X coordinate (Z = -23) of tunnel opening in SAG-SMO. A cluster is centered around the interface of TM2 and TM3, circled in red. The various helical boundaries are shown using black lines.

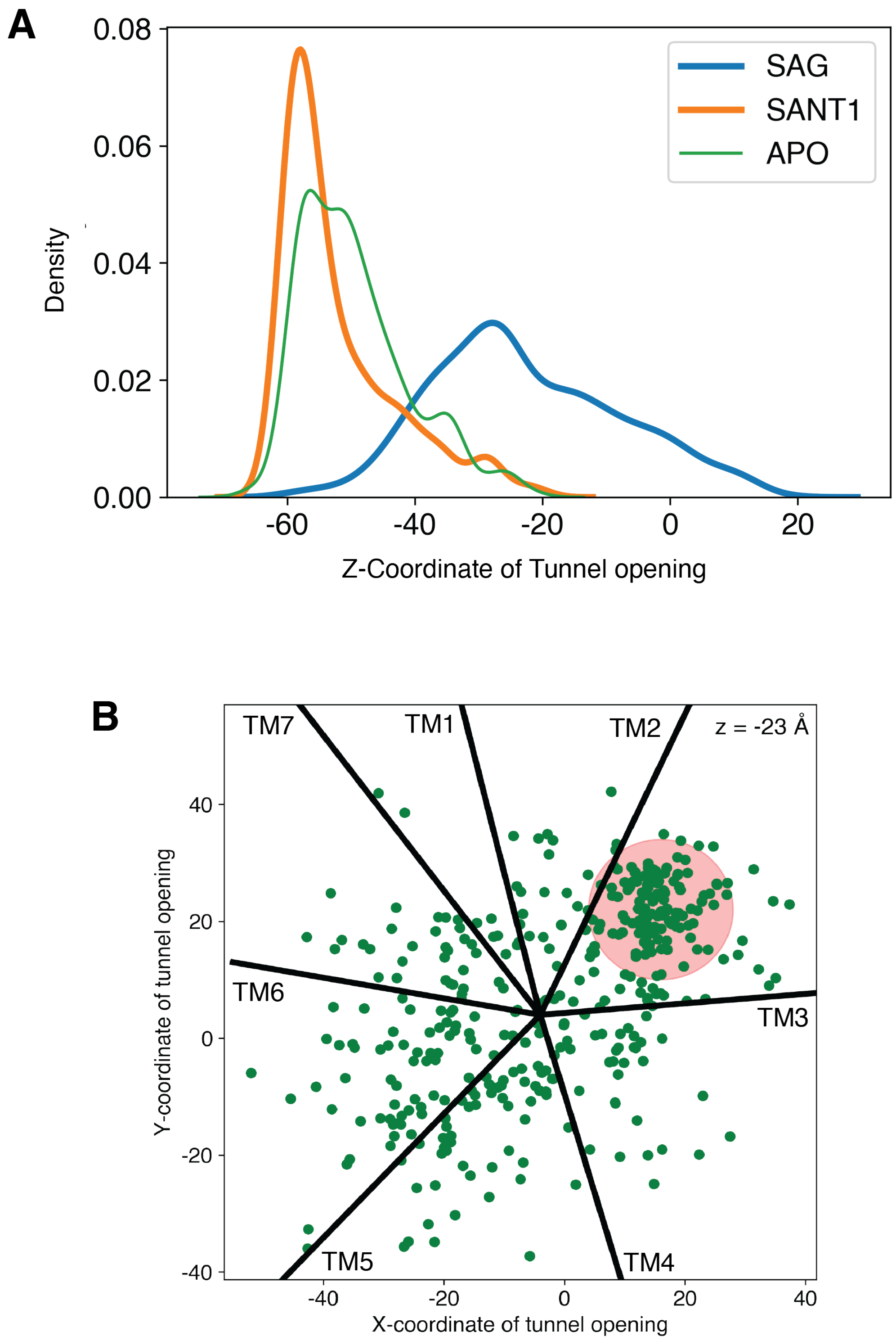


Fig. S24: (A) Probability density of Z-coordinate of tunnel opening. Simulations show that the tunnel opens in the upper leaflet of the membrane, z $\sim$ -22. (B) The Y v/s X coordinate (Z = -23) of tunnel opening in SAG-SMO. A cluster is centered around the interface of TM2 and TM3, circled in red. The various helical boundaries are shown using black lines.

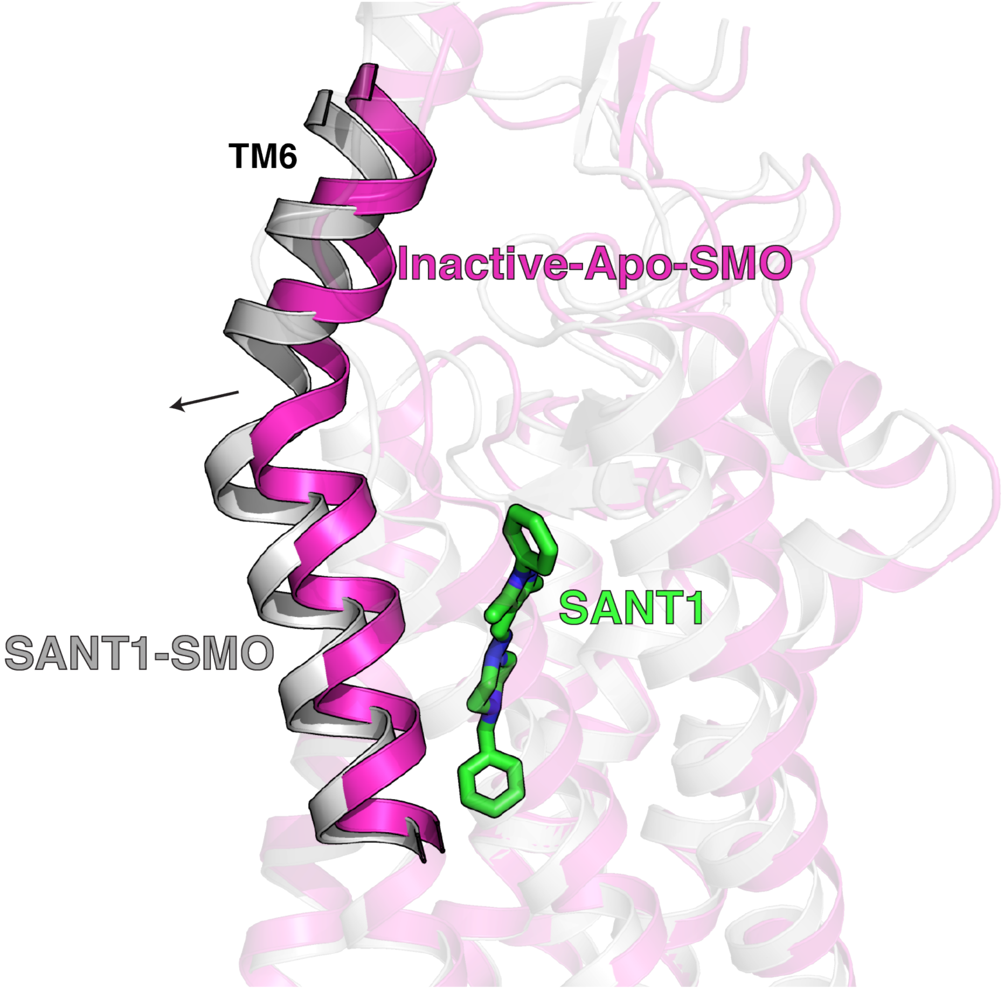


Fig. S25: Helical displacement of TM6 by SANT1. This helical displacement causes the shift in the allosteric network of SMO, precluding the transition to active state.

Table S1: Round wise data collection for Apo-SMO.

| Simulation Round | Amount of Apo-SMO-Data |
|---|---|
| Round 1 | 22$\mu$s |
| Round 2 | 23$\mu$s |
| Round 3 | 53$\mu$s |
| Round 4 | 46$\mu$s |
| Round 5 | 43$\mu$s |
| Round 6 | 40$\mu$s |
| Round 7 | 24$\mu$s |
| Total | 251 $\mu$s |

Table S2: Round wise data collection for SAG-SMO and SANT1-SMO

| Simulation Round | Amount of SAG-SMO-Data | Amount of SANT1-SMO-Data |
|---|---|---|
| Round 1 | 10$\mu$s | 10$\mu$s |
| Round 2 | 12$\mu$s | 14$\mu$s |
| Round 3 | 14$\mu$s | 18$\mu$s |
| Total | 36 $\mu$s | 42$\mu$s |

Table S3: Modelled residues in 5L7D-inactive-Apo-SMO starting structure. The helical content of K440-I445 was modelled based on the structure of SANT1-bound SMO (PDB: 4N4W)[8]

| Modelled Residues in 5L7D-inac-Apo-SMO | Constraints | Location |
|---|---|---|
| I429 | None | ICL3 |
| K430 | None | ICL3 |
| S431 | None | ICL3 |
| N432 | None | ICL3 |
| H433 | None | ICL3 |
| P434 | None | ICL3 |
| G435 | None | ICL3 |
| L436 | None | ICL3 |
| L437 | None | ICL3 |
| S438 | None | ICL3 |
| E439 | None | ICL3 |
| K440 | $\alpha$-helical | TM6 |
| A441 | $\alpha$-helical | TM6 |
| A442 | $\alpha$-helical | TM6 |
| S443 | $\alpha$-helical | TM6 |
| K444 | $\alpha$-helical | TM6 |
| I445 | $\alpha$-helical | TM6 |

Table S4: Membrane composition used for simulations.

| Lipid | Upper Leaflet | Lower Leaflet |
|---|---|---|
| Cholesterol | 21 | 21 |
| POPC | 76 | 76 |
| Sphingomyelin | 4 | 4 |
| Total | 101 | 101 |

Table S5: Adaptive Sampling metrics used for clustering in Apo-SMO, SANT1-SMO and SAG-SMO. Double lines separate pairs of residues.

| | | | | | |
|---|---|---|---|---|---|
| P58 | P59 | R66 | R173 | Y85 | K133 |
| L106 | Y130 | L108 | W109 | W109 | L126 |
| R113 | W119 | A115 | E211 | R117 | W206 |
| R117 | G212 | W119 | Q123 | Q123 | F187 |
| R151 | W163 | W163 | R168 | C169 | F174 |
| N202 | W206 | W206 | E211 | W206 | G212 |
| Y207 | E208 | Y207 | D209 | F222 | L515 |
| H231 | F285 | H231 | R290 | L246 | F275 |
| F252 | S259 | F252 | F268 | W256 | F268 |
| Y262 | L353 | Q284 | R290 | G288 | E292 |
| R290 | R291 | F332 | A459 | L335 | W339 |
| W339 | G422 | W339 | M449 | W339 | G453 |
| W339 | W535 | F343 | M449 | L346 | I445 |
| L353 | F360 | K356 | F360 | Q380 | Y399 |
| V381 | V392 | Y397 | F474 | Y397 | Q477 |
| L419 | F457 | H433 | P434 | H433 | G435 |
| G435 | L436 | S443 | N446 | H470 | F474 |
| W480 | P513 | Y487 | Q491 | Y487 | I509 |
| Q502 | I504 | L516 | K519 | T534 | W537 |
